# Entorhinal cortex glutamatergic and GABAergic projections bidirectionally control discrimination and generalization of hippocampal representations

**DOI:** 10.1101/2023.11.08.566107

**Authors:** Vincent Robert, Keelin O’Neil, Shannon K. Rashid, Cara D. Johnson, Rodrigo G. De La Torre, Boris V. Zemelman, Claudia Clopath, Jayeeta Basu

## Abstract

Discrimination and generalization are crucial brain-wide functions for memory and object recognition that utilize pattern separation and completion computations. Circuit mechanisms supporting these operations remain enigmatic. We show lateral entorhinal cortex glutamatergic (LEC_GLU_) and GABAergic (LEC_GABA_) projections are essential for object recognition memory. Silencing LEC_GLU_ during *in vivo* two-photon imaging increased the population of active CA3 pyramidal cells but decreased activity rates, suggesting a sparse coding function through local inhibition. Silencing LEC_GLU_ also decreased place cell remapping between different environments validating this circuit drives pattern separation and context discrimination. Optogenetic circuit mapping confirmed that LEC_GLU_ drives dominant feedforward inhibition to prevent CA3 somatic and dendritic spikes. However, conjunctively active LEC_GABA_ suppresses this local inhibition to disinhibit CA3 pyramidal neuron soma and selectively boost integrative output of LEC and CA3 recurrent network. LEC_GABA_ thus promotes pattern completion and context generalization. Indeed, without this disinhibitory input, CA3 place maps show decreased similarity between contexts. Our findings provide circuit mechanisms whereby long-range glutamatergic and GABAergic cortico-hippocampal inputs bidirectionally modulate pattern separation and completion, providing neuronal representations with a dynamic range for context discrimination and generalization.

## Introduction

Pattern separation and completion are neuronal computations that drive discrimination between dissimilar contexts while allowing for generalization across similar experiences to dynamically yield orthogonal and overlapping outputs, respectively^1–5^. Thereby, internal representations preserve their coherence but also adaptively remap given the contextual salience. While pattern separation and completion have been observed across the brain and modeled extensively^1–14^, the specific circuit mechanisms allowing for balanced pattern separation and completion operations within a given neural network remain enigmatic. In this study, we examined the specific roles of long-range and local circuit interactions in area CA3 to resolve how a single brain area can dynamically perform these seemingly opposing information processing operations.

As a model system akin to many cortical recurrent networks, hippocampal area CA3 has the unique ability to perform both pattern completion and separation^12,15–18^. Canonically, modeling and experimental work predict that attractor dynamics of the recurrently connected CA3 network promote pattern completion^1–3,6,8,9,12,13,19,20^, whereas the orthogonalization driven by the sparsely organized dentate gyrus (DG) inputs favors pattern separation^2,3,7,10–12,14,21–25^. However, CA3 also receives spatial and contextual inputs from the medial and lateral entorhinal cortex (MEC^26–29^ and LEC^30–38^). We know little about how these cortical inputs integrate with the local circuits and shape the crucial mnemonic computations underlying discrimination and generalization^39–41^. Within EC, LEC is a strong candidate to arbitrate the balance of pattern separation versus completion in area CA3^12,15–18^. Canonically, LEC is known for coding non-spatial contextual features of an environment^30^, including objects^31–34^, odors^35–37^, novelty^36^ and salient rewards^36,38^ or punishments^36^.

Further, LEC may also contribute to spatial coding given LEC lesions impair rate remapping of CA3 spatial representations^42^ (in-field firing change but not place field location), and paired extracellular recordings^30^ predict its involvement in pattern separation^12^ by reporting salient contextual changes. However, causal links and mechanistic insights can only be drawn with specific circuit manipulations to parse the little-known synaptic organization of long-range inputs from LEC to CA3 and their interactions with local circuits. Such information is crucial to understanding the cellular and network mechanisms underlying pattern separation and completion to support the formation and recall of neuronal representations across a range of context variations.

Long-range communication between LEC and CA3 relies not only on glutamatergic inputs (LEC_GLU_) but also on direct GABAergic projections (LEC_GABA_). Although better described in subcortical regions, studies exploring connectivity and function of long-range GABAergic projections across cortical areas^36,43–47^ are coming of age. Recent studies uncovered key roles for long-range GABAergic projections from LEC to hippocampal area CA1 in gating dendritic spikes, synaptic plasticity, and context discrimination^36^, as well as MEC to CA1 in modulating oscillatory synchronization^43^. However, it remains to be tested if and how LEC_GLU_ and LEC_GABA_ act conjunctively or independently to differentially gate information processing. Therefore, we investigated long-range glutamatergic and GABAergic circuit interactions between LEC and CA3 to gain insights into how molecularly-defined projections shape specialized computations underlying varied mnemonic functions distributed within the hippocampus^4,5^.

While area CA3 provides a major excitatory drive to the main hippocampal output area CA1, the cellular properties, circuit organization, input-output transformations, and mnemonic functions of the two sub-regions are distinct^15–18,48,49^. In contrast to CA1, CA3 pyramidal neurons (PNs) are much leakier integrators^50,51^ and receive stratified dendritic input from many different sources. These include (i) the direct inputs to distal dendrites from LEC and MEC conveying multisensory contextual and spatial information, (ii) the feedforward processed inputs from DG to proximal dendrites thought to provide orthogonalized information allowing CA3 to perform discrimination through pattern separation^2,3,7,10–12,14,21–25^, and (iii) the feedback inputs from within CA3/2 through recurrent connections (RC) upon medial and basal dendrites implicated in generalization through pattern completion^1–3,6,8,9,12,13,19,20^. We hypothesize that LEC GABAergic inputs differentially control the integration of LEC glutamatergic inputs with DG and RC inputs, thereby influencing input/output transformations and related computations in area CA3 differently from area CA1. To test this, we examined the role of LEC glutamatergic and GABAergic inputs to hippocampal area CA3 in shaping behavioral and neuronal output at the single neuron sub-cellular and ensemble network levels in novelty- and context-dependent object and spatial coding paradigms. For this, we employed a multidisciplinary approach using cell type specific-chemogenetic manipulations during freely moving behavior as well as head-fixed *in vivo* two-photon calcium imaging, *ex vivo* somatic and dendritic patch-clamp electrophysiology combined with dual-color optogenetic circuit mapping, and attractor-based computational modeling.

## Results

### LEC sends direct glutamatergic (LEC_GLU_) and GABAergic (LEC_GABA_) projections to area CA3

To map LEC_GLU_ and LEC_GABA_ projections to hippocampal area CA3 in the same animals, we co-expressed Chronos-GFP^52^ in excitatory and ChrimsonR-tdTomato^52^ in inhibitory neurons in LEC (Figure 1A). Fluorescently-labelled axonal fibers from both LEC_GLU_ (GFP+) and LEC_GABA_ (TdTomato+) projection neurons were observed in the distal dendritic layer (SLM) across the hippocampus including area CA3 (Figure 1B-C). In separate animals, injections with virus expressing ChR2^53^ showed similar innervation patterns (Figure S1A-B). All further experiments targeted the dorsal-most third of the hippocampus, typically associated with spatial memory^54,55^, hence providing an ideal substrate to probe pattern separation and completion.

**Figure 1.**
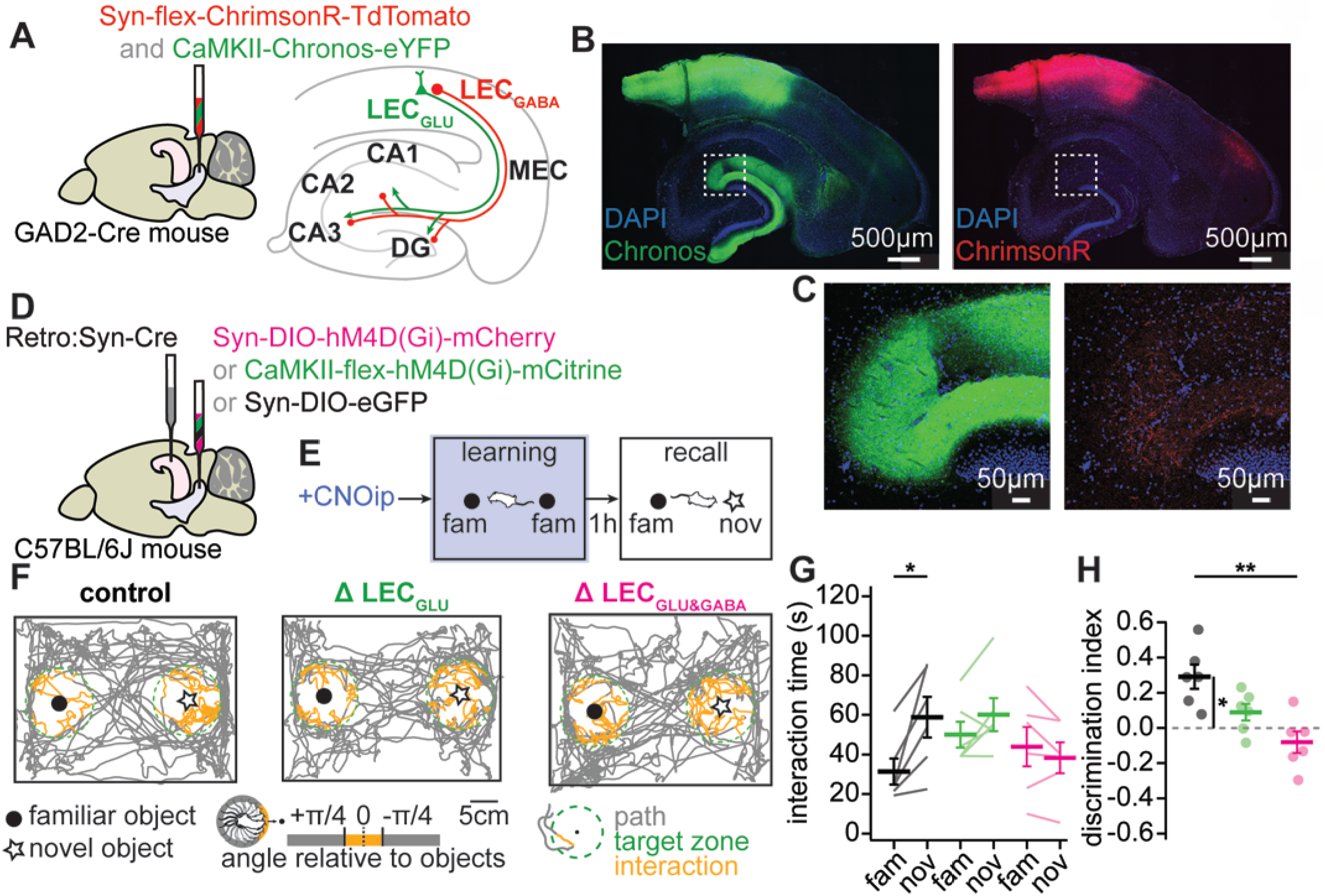
LEC_GLU_ and LEC_GABA_ inputs to CA3 jointly support object and cue discrimination. **(A)** Viral strategy. **(B)** Sample injection site of AAVs targeting LEC_GLU_ (GFP, green, left) and LEC_GABA_ (TdTomato, red, right) neurons and their projections to CA3, with DAPI staining (blue). **(C)** Expanded views of demarcated area in (B) showing LEC_GLU_ (left) and LEC_GABA_ (right) axons in CA3 SLM. **(D)** Viral strategy. **(E)** Experimental design. **(F)** Sample NOR recall sessions showing mouse path (grey) in the open field (black outline) and exploration of the familiar (closed circle) and novel (open circle) objects. Bouts of interaction with the objects (orange) are defined as the animals being within target zones (green dashed circles) centered around the objects and facing the corresponding object (head-body angle of mouse spanning +/− 45° relative to the object). **(G)** Object interaction time (control, n = 6, paired-T test, p = 0.018; Δ LEC_GLU_, n = 6, paired-T test, p = 0.114; Δ LEC_GLU&GABA_, n = 6, paired-T test, p = 0.205). **(H)** Discrimination index (control, n = 6, Δ LEC_GLU_, n = 6, Δ LEC_GLU&GABA_, n = 6, one-way ANOVA, p = 0.002, Tukey *post hoc* test, control vs Δ LEC_GLU_, p = 0.076, control vs Δ LEC_GLU&GABA_, p = 0.001, Δ LEC_GLU_ vs Δ LEC_GLU&GABA_, p = 0.146). Error bars represent SEM.

### Learning of object recognition memory requires LEC_GLU_ and LEC_GABA_ inputs to area CA3

To delineate the behavioral role of LEC_GLU_ and LEC_GABA_ inputs to area CA3, we chemogenetically silenced either LEC_GLU_ or LEC_GLU&GABA_ activity with Designer Receptors Exclusively Activated by Designer Drugs^56^ (DREADD, Figure S2A) in freely moving mice during hippocampal-dependent episodic memory tasks. Gross lesions of LEC affect object recognition memory^57–59^ and LEC neurons display object-related activity^31–34,60^. Therefore, we first tested the effect of pathway-specific silencing by clozapine N-oxide (CNO) injection during the learning phase of a novel object recognition (NOR) task in mice (Figure 1D-F, Figure S2B-D). Removal of LEC_GLU_ input to CA3 during the NOR encoding session decreased the novel object discrimination index during the recall session to chance levels, although this effect did not reach statistical significance compared to control (Figure 1G-H). Silencing both CA3-projecting LEC_GLU&GABA_ however significantly impaired performance (Figure 1G-H), suggesting that LEC_GLU_ and LEC_GABA_ may promote non-overlapping CA3 computation in favor of learning and memory. In contrast, spatial learning and recall performance in the Barnes maze was unaffected by LEC_GLU_ or LEC_GLU&GABA_ silencing (Figure S3), suggesting that LEC inputs are not required for spatial learning as assessed by this paradigm. Taken together, these observations indicate that LEC_GLU_ and LEC_GABA_ projections to CA3 are involved in object discrimination but not spatial learning. Further, the Δ LEC_GLU_ and Δ LEC_GLU&GABA_ groups counterintuitively showed overlapping phenotypes. Indeed, silencing of LEC_GLU&GABA_ tended to exacerbate the NOR impairment exhibited by the Δ LEC_GLU_ group, suggesting that LEC_GLU_ and LEC_GABA_ may be functionally complementary and therefore produce compounding rather than opposite effects when silenced. This prompted us to look further into the contributions of LEC_GLU_ and LEC_GABA_ to CA3 activity *in vivo*.

### LEC_GLU_ and LEC_GABA_ bidirectionally modulate CA3 PN population dynamics

To answer how LEC_GLU_ and LEC_GABA_ contribute to CA3 PN activity at the population level, we performed *in vivo* 2-photon calcium imaging of CA3 PN somas and apical dendrites in head-fixed animals running for water rewards on a double sided multi-textured Möbius belt (see methods) that allowed rapid switching between sides. We expressed Pharmacologically Selective Actuator Module (PSAM) ^36,61–63^ in LEC_GLU_ and DREADD in LEC_GABA_ to silence either the former alone (Δ LEC_GLU_) or both (Δ LEC_GLU&GABA_) and imaged calcium transients from GCaMP6f-expressing CA3 PNs (Figure 2A-D). After run training for randomly delivered water rewards (random foraging, RF) for 9 days on the same side of the belt (familiar context), mice were imaged with or without silencing of LEC_GLU_ or LEC_GLU&GABA_ in the familiar context, followed by exposure to the novel side of the belt (novel context).

**Figure 2.**
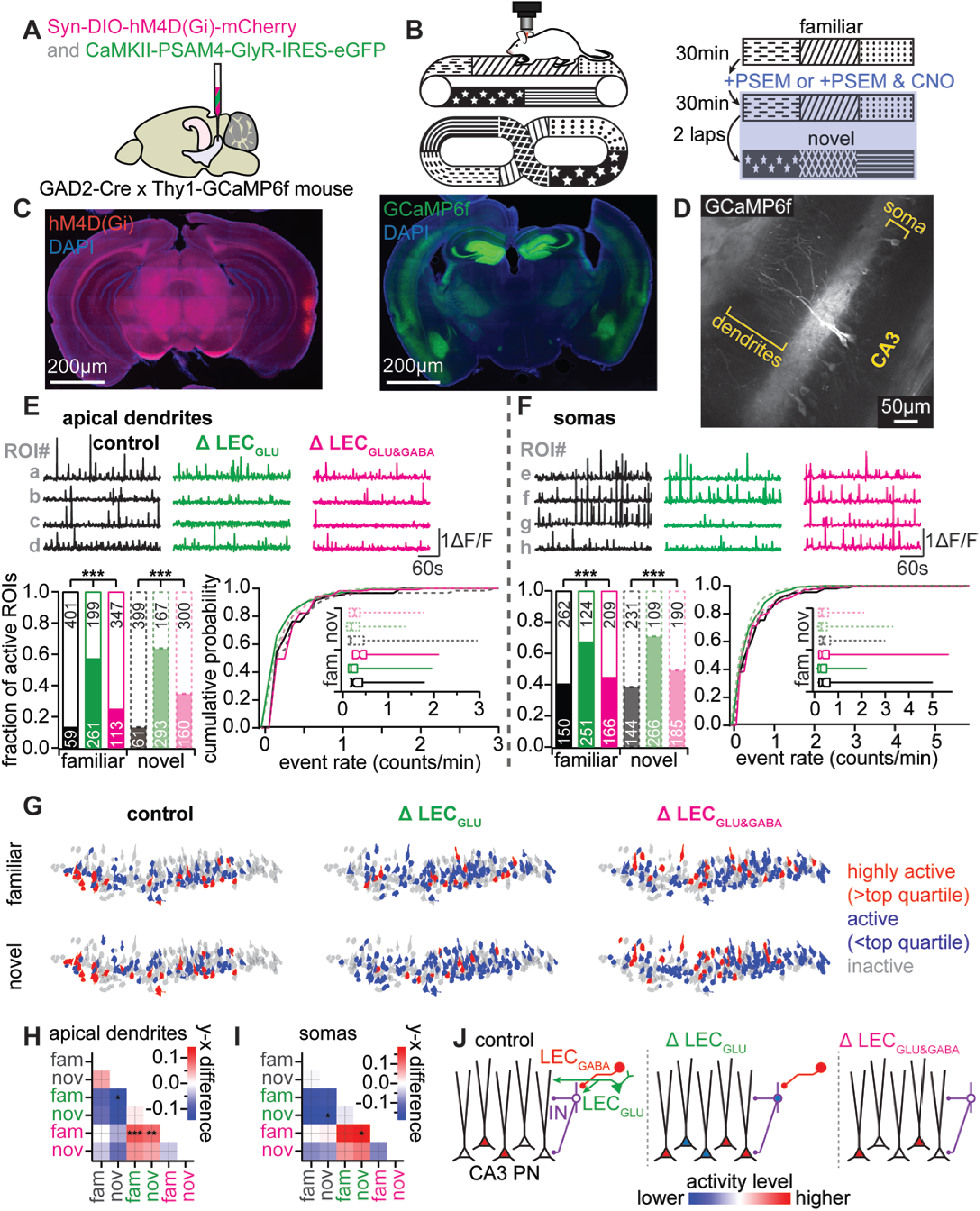
LEC_GLU_ and LEC_GABA_ modulate CA3 PN activity rate and sparseness. **(A)** Viral strategy. **(B)** Experimental design. **(C)** Left, sample injection site of AAVs targeting LEC_GLU_ and LEC_GABA_ (mCherry, red) neurons, with DAPI staining (blue). Right, sample imaging site of CA3 PNs (GCaMP6f, green), with DAPI staining (blue). **(D)** Sample CA3 field of view. **(E)** Sample traces (top), fraction of active ROIs (bottom, left; familiar: χ^2^ test, p < 0.001; novel: χ^2^ test, p < 0.001) and calcium transient rate (bottom, right; familiar: control, n = 59, Δ LEC_GLU_, n = 261, Δ LEC_GLU&GABA_, n = 113, Kruskal-Wallis ANOVA, p < 0.001; novel: control, n = 61, Δ LEC_GLU_, n = 293, Δ LEC_GLU&GABA_, n =160, Kruskal-Wallis ANOVA, p < 0.001) of CA3 PN apical dendrites during the familiar (solid lines) and novel (dashed lines) sessions in control (saline, black), Δ LEC_GLU_ (PSEM, green) and Δ LEC_GLU&GABA_ (PSEM & CNO, magenta) conditions. **(F)** Same as (E) but with CA3 PN somas (fraction of active ROIs, familiar: χ^2^ test, p < 0.001; novel: χ^2^ test, p < 0.001), (event rate, familiar: control, n =150, Δ LEC_GLU_, n = 251, Δ LEC_GLU&GABA_, n = 166, Kruskal-Wallis ANOVA, p < 0.001; novel: control, n =144, Δ LEC_GLU_, n = 266, Δ LEC_GLU&GABA_, n =185, Kruskal-Wallis ANOVA, p < 0.001). **(G)** Sample ROI footprints color-coded according to their activity rate. **(H)** Difference matrix of active CA3 PN dendrites event rates. **(I)** Difference matrix of active CA3 PN somas event rates. **(J)** Schematic interpretation.

We found that silencing of LEC_GLU_ increased the fraction of active CA3 PN somas and dendrites while decreasing their calcium transient rates in both the familiar and novel environments (Figure 2E-I, Table S1). Further silencing of LEC_GLU&GABA_ resulted in both the fraction of active ROIs and rate of calcium transients to reset similar to control (Figure 2E-I, Table S1). This indicates that ongoing activity of LEC_GLU_ drives CA3 PN activity in a given population while restricting the number of CA3 PNs active at any given time. This occurs possibly by directly exciting both CA3 PNs and interneurons (INs) that in turn provide local inhibition in a feedforward manner to suppress the activity of neighboring CA3 PNs. Conversely, LEC_GABA_ appears to counteract the LEC_GLU_ influence over CA3 PN population dynamics in this paradigm. We hypothesized this may stem from: (i) direct inhibition of CA3 PNs by LEC_GABA_ opposing LEC_GLU_-driven excitation, (ii) direct inhibition of CA3 INs by LEC_GABA_ reducing LEC_GLU_-driven feedforward inhibition thereby yielding overall disinhibition of CA3 PNs, or (iii) lateral inhibition of CA3-projecting LEC_GLU_ by LEC_GABA_ in LEC. Altogether, these results indicate that the influence of LEC_GLU_ and LEC_GABA_ over CA3 is not simply excitatory and inhibitory respectively, but may additionally involve counteracting forces of feedforward inhibition and disinhibition (Figure 2J). These may contribute to contrast generation in CA3 PN ensembles by influencing the balance of activity rate and sparseness of CA3 population dynamics (Figure 2J).

### Inhibition controls the recruitment of CA3 PNs by LEC_GLU_

To test whether our observed increase in CA3 population activity *in vivo* upon silencing LEC_GLU_ stemmed from recruitment of feedforward inhibition, we functionally mapped the LEC_GLU_ to CA3 connection using optogenetics and acute slice electrophysiology. We performed whole-cell patch-clamp recordings of CA3 PN responses evoked by photostimulation of Chronos or ChR2 expressing LEC_GLU_ inputs (Figure 3A), as well as electrical stimulations of local DG mossy fiber (MF, DCG-IV sensitive, Figure S4A) and CA3 recurrent collateral (RC) inputs (Figure 3B). This way we probed both pathway-specific and integrative properties of the major inputs to CA3 PNs.

**Figure 3.**
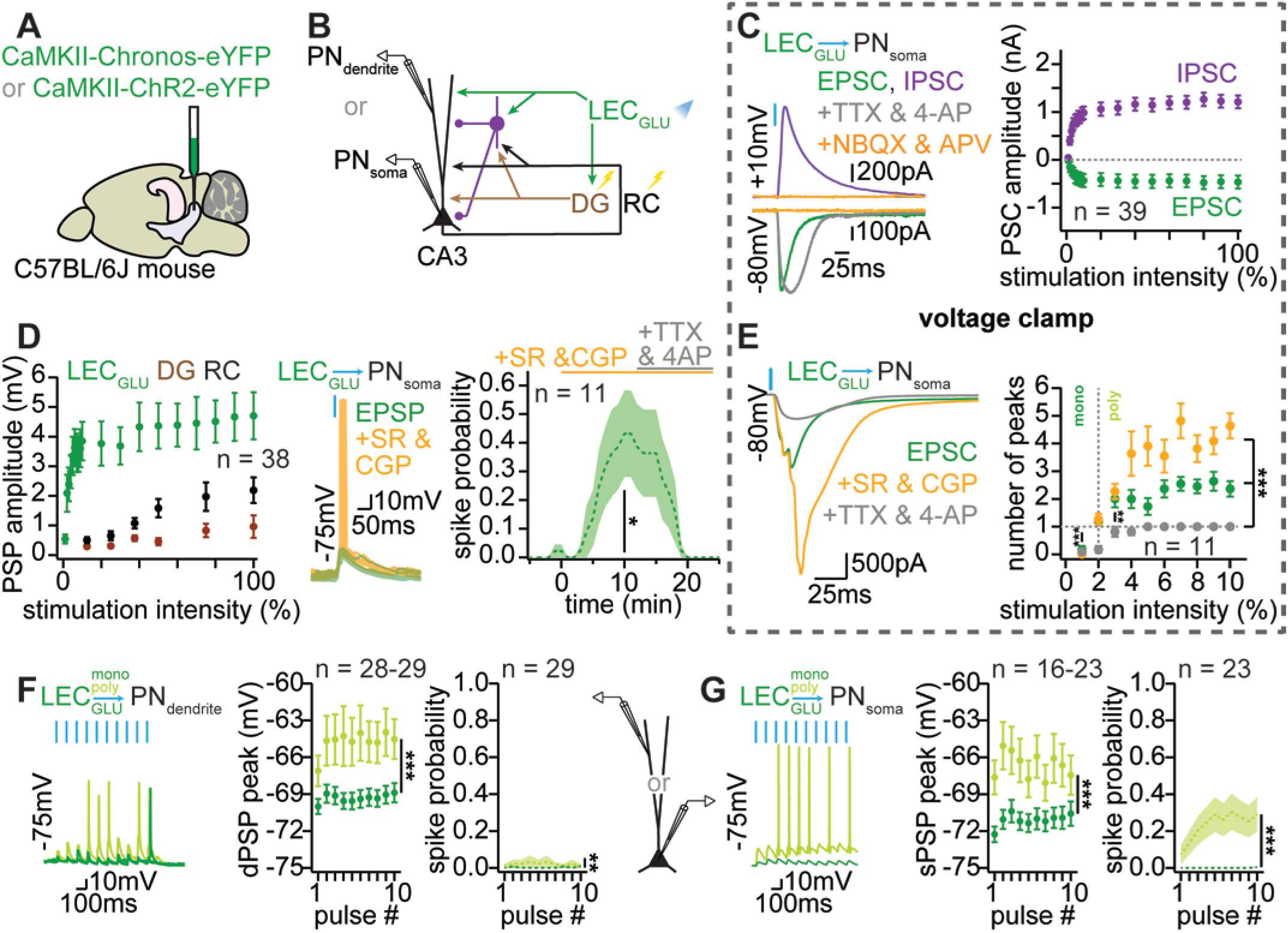
LEC_GLU_ drives excitation and feedforward inhibition in CA3 PNs. **(A)** Viral strategy. **(B)** Experimental design. **(C)** Left, sample traces of LEC_GLU_-driven CA3 PN post-synaptic responses in ACSF (EPSC, green; IPSC, purple), TTX & 4-AP (grey), and NBQX & APV (orange). Right, input-output curves of LEC_GLU_-evoked CA3 PN EPSC (green) and IPSC (purple) amplitudes. **(D)** Left, input-output curves of LEC_GLU_-(green), DG-(brown), and RC-evoked (black) CA3 PN PSP amplitudes. Middle, sample traces of LEC_GLU_-evoked CA3 PN PSPs and spikes in ACSF (green) and SR & CGP (orange). Right, time-course of LEC_GLU_-evoked CA3 PN spike probability (spike probability in SR & CGP: n = 11, one-sample T test versus hypothetical mean of zero, p = 0.024). **(E)** Left, sample traces of LEC_GLU_-evoked CA3 PN EPSCs in ACSF (green), SR & CGP (orange), and TTX & 4-AP (grey). Right, input-output curves of the number of peaks in the LEC_GLU_-evoked CA3 PN EPSC waveform in ACSF (green), SR & CGP (orange), and TTX & 4-AP (grey) (number of EPSC peaks: n = 11, two-way ANOVA, treatment, p < 0.001, power, p < 0.001, treatment x power, p < 0.001), (number of EPSC peaks against LED power: n = 11, one-sample T tests versus hypothetical mean of one: 1 %, 0.07 mW/mm^2^, p < 0.001, 2 %, 0.14 mW/mm^2^, p = 0.167, 3 %, 0.21 mW/mm^2^, p = 0.008). **(F)** Sample traces, PSP peak (closed circles) and spike probability (dashed line) of CA3 PN dendritic responses to repeated stimulation of LEC_GLU_ monosynaptic or polysynaptic input at 10 Hz (PSP peak: n = 28 out of 29 samples because spikes were excluded from measurements, two-way ANOVA, polysynaptic vs monosynaptic, p < 0.001, pulse #, p = 0.971, polysynaptic vs monosynaptic x pulse #, p = 0.999; spike probability: n = 29, two-way ANOVA, polysynaptic vs monosynaptic, p = 0.008, pulse #, p > 0.999, polysynaptic vs monosynaptic x pulse #, p = 0.991). **(G)** Same as (F) but with CA3 PN somatic responses (PSP peak: n = 16 out of 23 samples because spikes were excluded from measurements, two-way ANOVA, polysynaptic vs monosynaptic, p < 0.001, pulse #, p = 0.825, polysynaptic vs monosynaptic x pulse #, p = 0.989; spike probability: n = 23, two-way ANOVA, polysynaptic vs monosynaptic, p < 0.001, pulse #, p = 0.618, polysynaptic vs monosynaptic x pulse #, p = 0.633). Error bars represent SEM.

Voltage-clamp recordings from CA3 PNs revealed that LEC_GLU_ stimulation evoked excitatory post-synaptic currents (EPSCs) and inhibitory post-synaptic currents (IPSCs) (Figure 3C). Application of the AMPAR and NMDAR receptor antagonists NBQX and APV abolished both EPSCs and IPSCs, showing that the LEC_GLU_ to CA3 PN transmission is purely glutamatergic and suggesting that EPSCs may be direct responses. In contrast, IPSCs may result from feedforward inhibition (Figure 3C, Figure S4B). This was confirmed by blocking all polysynaptic responses with the application of TTX and 4-AP, which spared LEC_GLU_-driven EPSCs but abolished IPSCs (Figure 3C, Figure S4C). Hence, LEC_GLU_ inputs drive monosynaptic glutamatergic excitation and disynaptic feedforward inhibition onto CA3 PNs. We next asked how these compound excitatory and inhibitory drives would affect CA3 PN membrane potential (V_M_) by sampling input-output curves while holding CA3 PNs at −70 mV near their resting V_M_ under current-clamp conditions (Figure 3D). Although post-synaptic potentials (PSPs) increased with stronger input stimulation, CA3 PNs did not fire action potentials in response to any given single input stimulation (Figure 3D). This indicates that, in our experimental conditions, CA3 PN output may require decreased inhibition and / or combined excitation from multiple inputs.

To test whether the feedforward inhibition recruited by LEC _GLU_ effectively curtails excitability in CA3 PNs, we compared PSP and spike output with current-clamp recordings before and after inhibition blockade with GABA_A_R and GABA_B_R antagonists (SR95531, SR and CGP55845A, CGP, respectively). While all responses with GABAergic transmission intact remained subthreshold, blockade of GABAergic transmission unmasked/ enabled action potential firing in CA3 PN soma (Figure 3D). This could stem from a relief of CA3 PN from shunting inhibition as well as the recruitment of additional excitation in the hippocampal network. To test the latter, we sampled CA3 PNs EPSC input-output curves in response to LEC_GLU_ stimulation first in ACSF as a reference, then in SR & CGP to gauge the maximal recruitment of excitation, and finally in TTX & 4-AP to confirm monosynaptic connection (Figure 3E, Figure S4E-G). Resolving the number of peaks in the EPSC waveforms, we found that LEC_GLU_-evoked EPSCs displayed more multiple peaks with stimulation intensity and were more numerous with GABAergic transmission blocked (Figure 3E). Further, multipeak EPSCs could be evoked with optogenetic stimulation of LEC_GLU_ above 3 % (0.21 mW/mm^2^) LED intensity in regular ACSF (Figure 3E), thus suggesting that strong LEC_GLU_ stimulation can recruit polysynaptic excitation from DG and/or CA3/2 by LEC_GLU_ inputs. We confirmed this by increasing LEC_GLU_ stimulation intensity while performing extracellular field potential recordings in DG GC and CA3 SP layers which readily displayed population spikes when 470 nm light power exceeded 5 % (0.33 mW/mm^2^) (Figure S4H-J), thus indicating that LEC_GLU_ can indeed recruit DG and RC as additional sources of excitation onto CA3 PNs. Conversely, these observations suggest that monosynaptic LEC_GLU_ input could be probed below our system’s 3 % (0.21 mW/mm2) LED power threshold (Figure 3E).

Quantification of the monosynaptic and polysynaptic LEC_GLU_, DG, and RC inputs onto CA3 PNs revealed substantial excitatory and inhibitory drives (Figure S4K-L). This led to significant compartment-specific depolarizations that remain subthreshold when inputs are stimulated in isolation with GABAergic transmission intact (Figure S4M-N). Altogether, these findings suggest that the dynamics of input integration and relief from feedforward inhibition may control spike output in CA3 PNs.

### LEC_GLU_ can recruit DG and RC local inputs to yield CA3 PN output

Since no single input could drive firing in CA3 PN, but LEC_GLU_ may polysynaptically recruit local DG and RC inputs, we asked whether repeated or combined inputs elicit CA3 PN spiking with *ex vivo* electrophysiology. First, we repeatedly stimulated LEC_GLU_ either at low (2 %, 0.14 mW/mm^2^, monosynaptic) or high (100 %, 3.76 mW/mm^2^, polysynaptic) light intensity while recording PSPs and spikes from CA3 PNs soma or dendrites (Figure 3F-G). The low stimulation intensity regime yielded small depolarizations and virtually no spiking (Figure 3F-G), consistent with activation of the monosynaptic LEC_GLU_-CA3 transmission, which targets the distal dendrites of CA3 PNs and therefore generates PSPs that attenuate before reaching the soma. In contrast, stimulating LEC_GLU_ at high power induced PSP summation and spiking during the train (Figure 3F-G) suggesting that recruitment of DG- and/or RC-driven polysynaptic excitation by LEC_GLU_ could drive CA3 dendritic spiking and somatic output.

To confirm that, we next probed whether high intensity (100 %, 3.76 mW/mm^2^) stimulation of LEC_GLU_ yielding polysynaptic excitation would occlude some of the additive effects of pairing LEC_GLU_ with DG or RC. For this, we delivered high intensity repeated stimulations of LEC_GLU_, DG, RC either alone or combined with one another while monitoring PSPs and spikes in CA3 PN somas or dendrites. First, we observed that each individual pathway repeated stimulation could elicit modest spiking in post-synaptic CA3 PNs (Figure S5A-D). Next, we found that combining DG and LEC_GLU_ high power stimulation did not yield significantly more spikes than expected from the linear arithmetic sum of the two inputs alone (Figure S5A, S5C). Similar results were obtained with LEC_GLU_ and RC pathways (Figure S5B, S5D). Thus, these results reveal an overlap between the polysynaptic LEC_GLU_ and the DG or RC pathways, as expected if high intensity LEC_GLU_ stimulation partially recruits DG or RC to drive polysynaptic excitation onto CA3 PN. Therefore, the ceiling effect of CA3 output in response to combined polysynaptic LEC_GLU_ and DG or RC suggests that the LEC_GLU_ input can physiologically recruit both DG and RC local inputs to drive activity in CA3 PNs.

### LEC_GABA_ activation decreases feedforward inhibition in CA3 PNs

Given we established that inhibition is a crucial determinant of LEC_GLU_ influence over CA3 PN activity, we next asked whether LEC_GABA_ could modulate feedforward inhibition. Indeed, LEC_GABA_ have previously been reported to have a disinhibitory action over PN dendritic activity in CA1^36^. Thus, we expressed Chronos in LEC_GLU_ and ChrimsonR in LEC_GABA_ to simultaneously probe these inputs to CA3 PNs and INs at the single cell level using *ex vivo* electrophysiology (Figure 1A-C, 4A, Figure S6).

First, we found that a fraction of CA3 SLM INs received convergent input from both LEC_GLU_ and LEC_GABA_, using TTX & 4-AP application to confirm monosynaptic connectivity (Figure 4B-C, Figure S7A). As expected, LEC_GLU_-evoked EPSCs were glutamatergic whereas LEC_GABA_-evoked IPSCs were GABAergic, as they were respectively blocked by NBQX & APV and SR & CGP (Figure 4B-C, Figure S7B). Although most CA3 INs received LEC_GLU_ input, only a fraction of them was targeted by LEC_GABA_ input (Figure 4D). Further, we observed that LEC_GABA_-connected CA3 SLM INs received strong LEC_GLU_ and RC excitatory input but only weak excitation from DG (Figure S7C), which suggests that the LEC LEC_GABA_ disinhibitory action may selective to LEC_GLU_ and RC but not DG inputs. Next, we resolved which genetically-identified CA3 IN subpopulations received LEC_GABA_ input and found selective innervation of VIP (8/30) and CCK (1/34) INs but not PV (0/22) and SST (0/30) INs (Figure 4E, Figure S7D-I). These findings raise the possibility that LEC_GABA_ innervates a subpopulation of basket INs co-expressing VIP- and CCK, which have been described in CA1 to have somatic location in SLM and perisomatic axonal projections^64,65^. This suggests that LEC_GABA_ may exert a somatic compartment-specific disinhibitory effect on CA3 PNs. Finally, we monitored CA3 SLM IN spike output to LEC_GLU_ +/− LEC_GABA_ input stimulation with dual optogenetics in cell-attached mode and found that LEC_GABA_ indeed decreased action potential firing to LEC_GLU_ input in CA3 SLM INs (Figure 4F). This suggests that LEC_GABA_ may have a net disinhibitory effect onto CA3 PNs by reducing feedforward inhibition.

**Figure 4.**
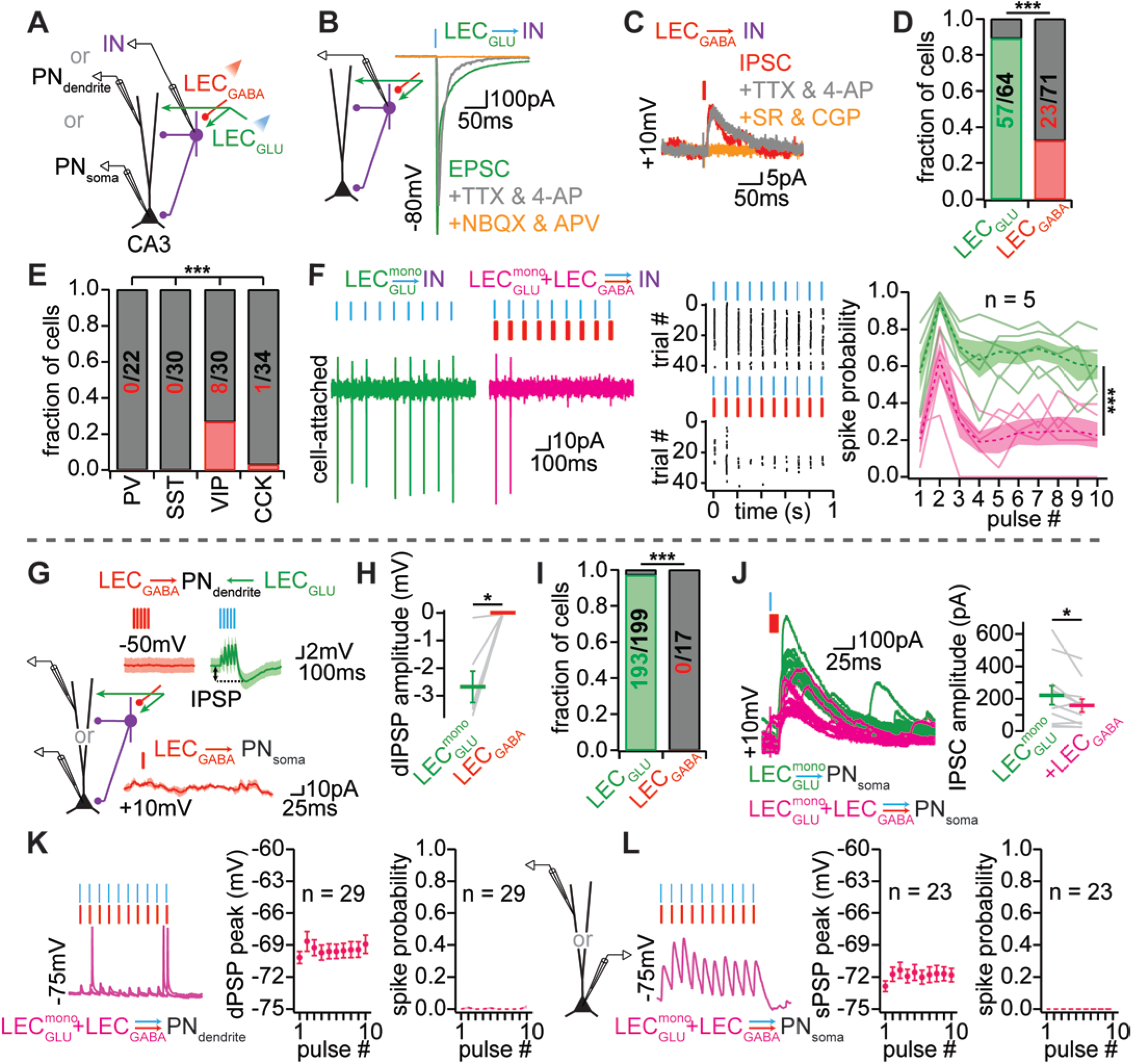
LEC_GABA_ reduces feedforward inhibition onto CA3 PNs. **(A)** Experimental design. **(B)** Sample traces of LEC_GLU_-evoked CA3 IN EPSCs in ACSF (green), TTX & 4-AP (grey), and NBQX & APV (orange). **(C)** Sample traces of LEC_GABA_-evoked CA3 IN IPSCs in ACSF (red), TTX & 4-AP (grey), and SR & GCP (orange). **(D)** Fraction of CA3 IN receiving LEC_GLU_ and LEC_GABA_ inputs. **(E)** Fraction of PV, SST, VIP and CCK INs receiving LEC_GABA_ input (χ^2^ test, p < 0.001). **(F)** Left, sample traces of CA3 IN action potential firing evoked by LEC_GLU_ (green) or LEC_GLU&GABA_ stimulation (magenta). Middle, sample raster plot. Right, spike probability of CA3 IN evoked by LEC_GLU_ with or without LEC_GABA_ stimulation (n = 5, two-way ANOVA, input combination, p < 0.001, pulse #, p < 0.001, input combination x pulse #, p = 0.982). **(G)** Top, sample traces of CA3 PN dendrites held at −50 mV under current-clamp upon LEC_GABA_ stimulation (left) and LEC_GLU_ stimulation (right), note the absence of hyperpolarization in response to LEC_GABA_ stimulation as opposed to feedforward inhibition-mediated hyperpolarization with LEC_GLU_ stimulation in the same samples. Bottom, sample traces of CA3 PN somas under voltage-clamp at +10 mV upon LEC_GABA_ stimulation. **(H)** CA3 PN dendritic IPSP amplitude evoked by stimulation of LEC_GLU_ and LEC_GABA_ (n = 6, Wilcoxon signed-rank test, p = 0.031). **(I)** Fraction of CA3 PN receiving LEC_GLU_ and LEC_GABA_ inputs. **(J)** Sample traces (left) amplitude (right) of LEC_GLU_-evoked CA3 PN IPSCs with (magenta) or without (green) LEC_GABA_ stimulation (n = 11, paired-T test, p = 0.042). **(K)** Sample traces, PSP peak (closed circles) and spike probability (dashed line) of CA3 PN dendritic responses to repeated stimulation of LEC_GLU_+LEC_GABA_ inputs at 10 Hz (comparison with corresponding LEC_GLU_ input presented in Figure 3F; PSP peak: n = 29, two-way ANOVA against LEC_GLU_ monosynaptic data, input combination, p = 0.578, pulse #, p = 0.893, input combination x pulse #, p > 0.999; spike probability: n = 29, two-way ANOVA, input combination, p = 0.670, pulse #, p = 0.081, input combination x pulse #, p = 0.947). **(L)** Same as (K) but with CA3 PN somatic responses (comparison with corresponding LEC_GLU_ input presented in Figure 3G; PSP amplitude: n = 23, two-way ANOVA, input combination, p = 0.120, pulse #, p = 0.759, input combination x pulse #, p > 0.999; spike probability: n = 23, two-way ANOVA, input combination, p = 0.318, pulse #, p = 0.439, input combination x pulse #, p = 0.439). Error bars represent SEM.

Next, we sought to establish whether LEC_GABA_ provided direct input to CA3 PNs, since the LEC_GABA_-hippocampal PN connection had never been probed before. In contrast to their LEC_GLU_ counterparts, photostimulation of ChrimsonR-expressing LEC_GABA_ failed to elicit post-synaptic responses of any kind in CA3 PNs, as evidenced by (i) the lack of time-locked IPSCs evoked by LEC_GABA_ stimulation in CA3 PN voltage-clamped at +10 mV (Figure 4G), (ii) the absence of LEC_GABA_-driven hyperpolarization in CA3 PN dendrites held at −50 mV where LEC_GLU_-driven feedforward inhibition results in V_M_ hyperpolarization (Figure 4G-H), and (iii) no GABA_A_R- and GABA_B_R-sensitive inhibitory field post-synaptic potential in response to LEC_GABA_ stimulation (Figure S7J); overall suggesting an absence of direct input from LEC_GABA_ to CA3 PNs (Figure 4G-I). Further, comparing the amplitude of LEC_GABA_-evoked IPSC (or lack thereof) in CA3 INs and PNs at the population level, we confirmed that CA3 INs are the sole target of LEC_GABA_ (Figure S7K). Indeed, while LEC_GLU_ innervation was near ubiquitous in both PNs and INs, LEC_GABA_ input was restricted to CA3 INs (Figure 4D, 4I). Finally, we tested our prediction that LEC_GABA_ input would have a disinhibitory effect by monitoring LEC_GLU_-, DG-, and RC-driven IPSCs in CA3 PNs with or without coincident activation of LEC_GABA_. We found that LEC_GABA_ decreased LEC_GLU_-driven feedforward inhibition onto CA3 PNs, thus indicating an input-specific disinhibitory mechanism (Figure 4J). In contrast, both DG-driven inhibition and RC-driven inhibition were unaffected by LEC_GABA_ stimulation (Figure S7L-M). These results predict that LEC_GABA_ may promote excitation by reducing LEC_GLU_-driven feedforward inhibition in CA3 PNs.

### LEC_GABA_ selectively boosts CA3 PN somatic output through input-specific disinhibition

To test whether the LEC_GABA_-mediated reduction of LEC_GLU_-driven feedforward inhibition disinhibits CA3 PN compartment-specific activity, we recorded CA3 PN somas or dendrites *ex vivo* and delivered repeated stimulations of LEC_GLU_ at low (2 %, 0.14 mW/mm^2^) light intensity either alone or combined with LEC_GABA_ (Figure 4K-L). Surprisingly, we found no significant difference in the PSP amplitude and spike probability in response to LEC_GLU_ with or without LEC_GABA_ in both somas and dendrites (Figure 4K-L). This suggests that, although LEC_GABA_ dampens LEC_GLU_-driven feedforward inhibition, the disinhibitory effect does not contribute to increasing CA3 PN post-synaptic responses to LEC_GLU_ input stimulation alone.

Since LEC_GABA_-driven disinhibition did not overtly change CA3 PN responses to LEC_GLU_ activation alone, we next asked whether the LEC_GABA_ disinhibitory action would promote integration of multiple inputs to yield CA3 PN output by virtue of gating pathway specific feedforward inhibition. For this, we tested if coincident stimulation of LEC_GABA_ with LEC_GLU_ and combinations of convergent DG or RC inputs would supralinearly increase PSPs and spike output in CA3 PN soma or dendrites *ex vivo* (Figure 5). We compared PSP peak and spike probability evoked by repeatedly stimulating LEC_GLU_, DG, and RC either alone or combined with one another, with or without additional LEC_GABA_ activation. We found that combined stimulations of LEC_GABA_ with LEC_GLU_ and RC together yielded more action potential firing in CA3 PN somas than expected by linear summation of LEC_GLU_ plus RC (Figure 5A-B). In contrast, similar pairings of LEC_GLU_ and DG with added stimulation of LEC_GABA_ did not show a supralinear increase in somatic action potential firing compared to their linear sum (Figure 5C-D). Because non-linearities in input integration can stem from dendritic spikes that help dendritic depolarizations from distal loci such as LEC_GLU_ reach the soma, we performed similar analysis of post-synaptic responses to LEC LEC_GABA_ pairing with LEC_GLU_ + DG or LEC_GLU_ + RC in our separate dendritic recording dataset. Surprisingly, although we sometimes observed dendritic spikes in response to LEC_GLU_ and RC pairing, additional stimulation of LEC_GABA_ did not significantly change dendritic spike probability (Figure 5E). Consistent with our somatic data, LEC_GLU_ and DG inputs did not sum efficiently in CA3 PN dendrites thus virtually never yielding dendritic spikes, and LEC_GABA_ had no further effect (Figure 5F). This suggests that LEC_GABA_ selectively boosts LEC_GLU_ integration with RC through a perisomatic disinhibitory mechanism.

**Figure 5.**
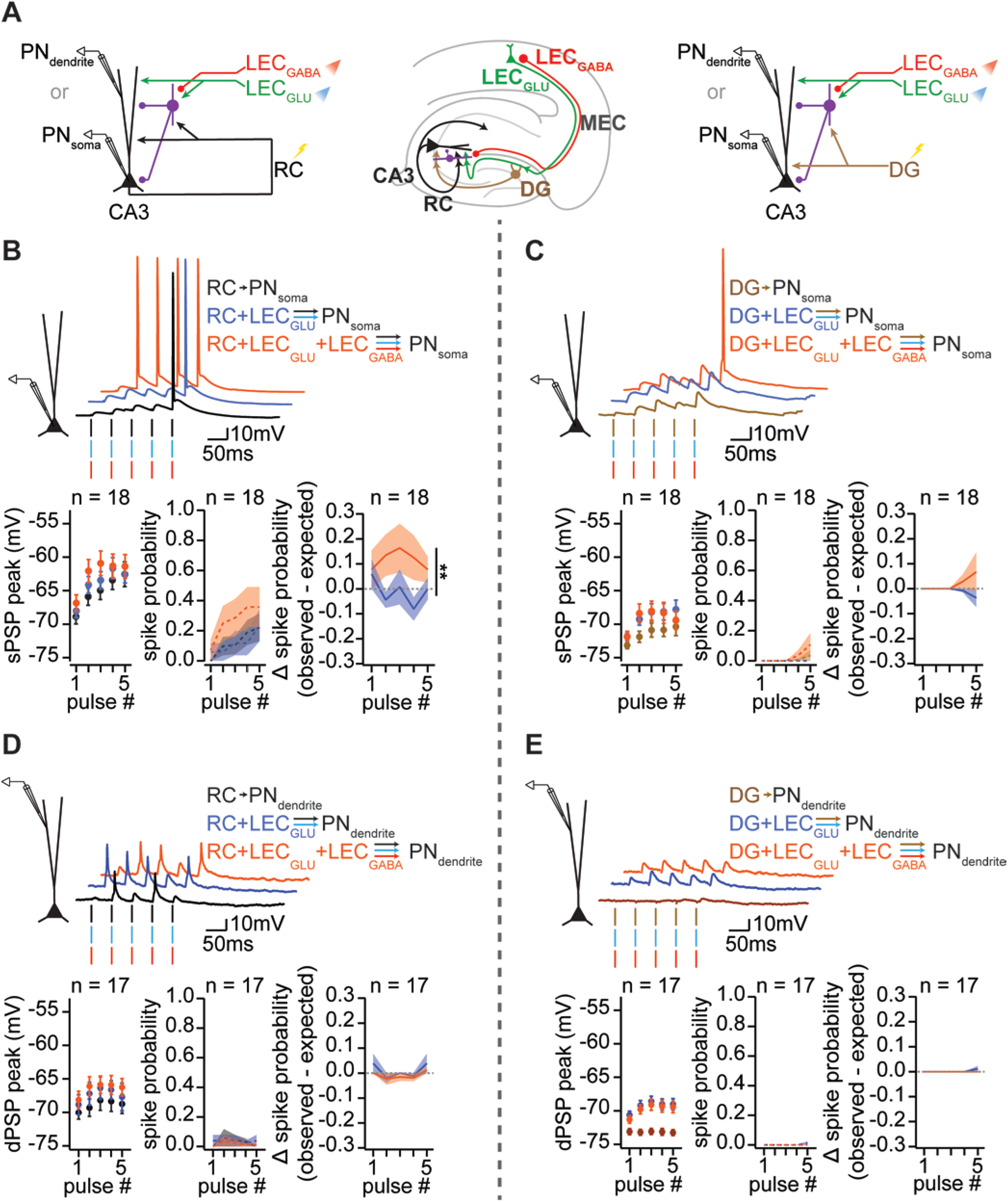
Input- and compartment-specific disinhibition from LEC inhibitory inputs boosts CA3 PN output. **(A)** Experimental design. **(B)** Sample traces (top), PSP peak and spike probability (bottom left) of LEC_GLU_-(green), RC-(black), LEC_GLU_ & LEC_GABA_-(magenta), LEC_GLU_ + RC-(blue), and LEC_GLU&GABA_ + RC-evoked (orange) CA3 PN somatic responses, and delta spike probability (bottom right) between observed RC + LEC_GLU_ with or without LEC_GABA_ stimulation versus expected from linear sum of these inputs (difference between observed vs expected spike probability: n = 18, two-way ANOVA, input combination, p = 0.007, pulse #, p = 0.841, input combination x pulse #, p = 0.549). **(C)** Experimental design. **(D)** Same as (B) with DG stimulation (brown) instead of RC (difference between observed vs expected spike probability: n = 18, two-way ANOVA, input combination, p = 0.115, pulse #, p = 0.993, input combination x pulse #, p = 0.258). **(E)** same as (B) but in CA3 PN dendrites (difference between observed vs expected spike probability, n = 17, two-way ANOVA, input combination, p = 0.173, pulse #, p = 0.111, input combination x pulse #, p = 0.925). **(F)** same as (D) but in CA3 PN dendrites (difference between observed vs expected spike probability, n = 17, two-way ANOVA, input combination, p = 0.319, pulse #, p = 0.409, input combination x pulse #, p = 0.409). Error bars represent SEM.

### LEC inputs contribute to CA3 remapping

Since LEC_GABA_ preferentially boosted integration of LEC_GLU_ with RC which is postulated to promote pattern completion but not DG inputs associated with pattern separation, we then examined their contributions to contextual change-induced remapping of CA3 place cells. We used the relative change of place field location at the population level as a proxy for pattern separation and completion. By favoring conjunctive LECGLU- and RC-driven activity in CA3, we hypothesized that LEC_GABA_ may tip the balance of remapping in CA3 ensembles towards pattern completion through increased generalization between distinct contexts conveyed by LEC_GLU_. In this framework, silencing the LEC_GLU_ inputs alone would result in accentuated pattern completion, yielding more similar spatial outputs even with different contextual inputs. Conversely, without both LEC_GABA_ and LEC_GLU_, CA3 representations may lose the dynamic range of remapping brought about by LEC inputs. To test this hypothesis, we chemogenetically silenced LEC_GLU_ alone or LEC_GLU&GABA_ together as mice performed a navigational goal-oriented learning (GOL) task with varying similarities between sensory cues, creating three different contexts (familiar, intermediate and novel). In this paradigm, head-fixed mice have to travel to a specific location on the belt to receive water rewards based on the sensory context^66,67^ while GCaMP6f-expressing CA3 PN somatic and dendritic activity is captured with two-photon *in vivo* calcium imaging. LEC_GLU_ (Δ LEC_GLU_) or LEC_GLU&GABA_ (Δ LEC_GLU&GABA_) were silenced as mice ran in familiar, intermediate (same tactile cues, different olfactive and auditory cues), and novel (different tactile, olfactive, and auditory cues) contexts (Figure 6A-B, Figure S8A).

**Figure 6.**
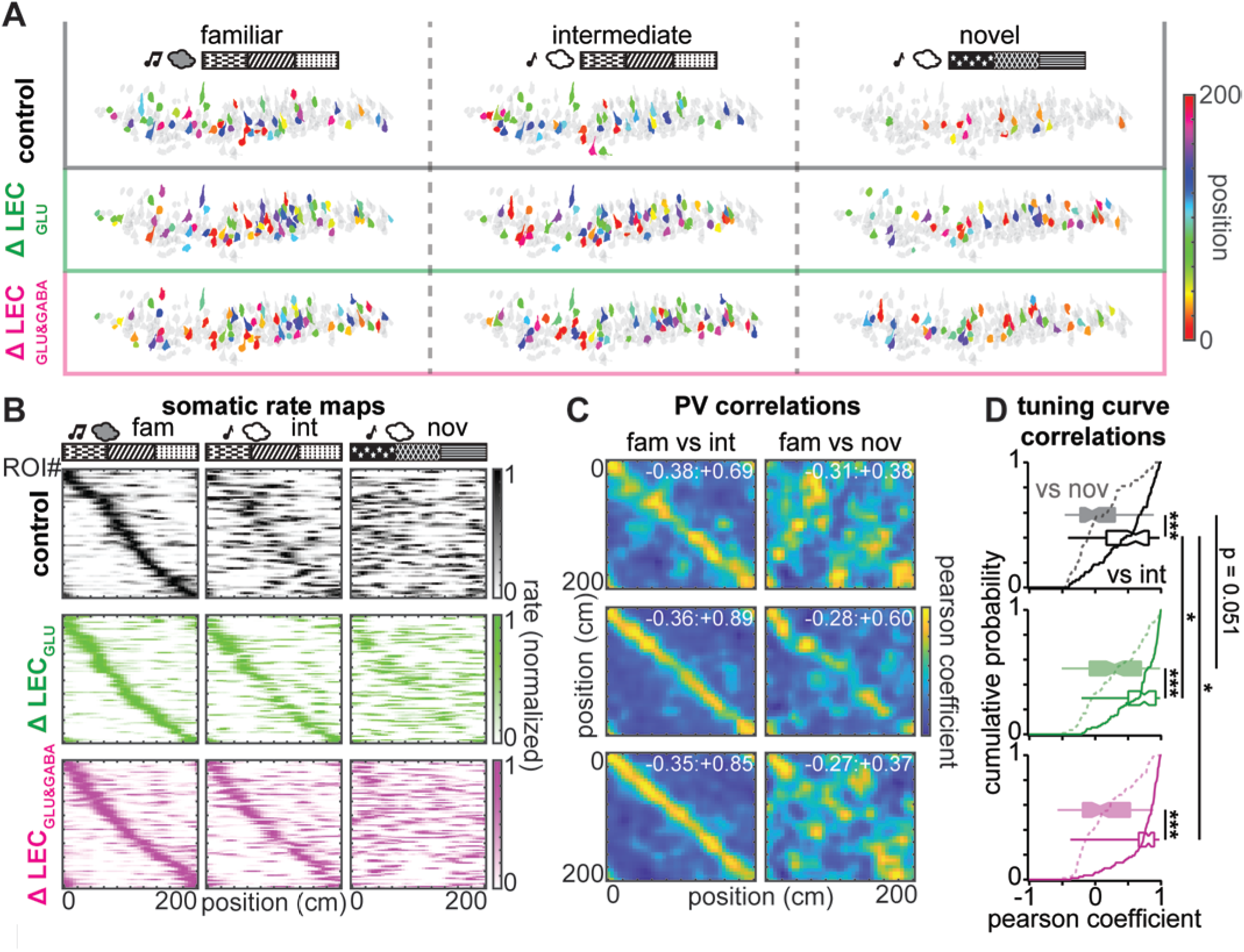
LEC_GLU_ & LEC_GABA_ bidirectionally modulate CA3 place cell remapping. **(A)** Sample ROI footprints color-coded according to the position of their place field. **(B)** Normalized rate maps of CA3 place cell somas sorted according to their place field location in the familiar session. Left to right: familiar, intermediate, and novel sessions. Top to bottom: control (black), Δ LEC_GLU_ (green), Δ LEC_GLU&GABA_ (magenta). Topmost schematics represent experimental design. **(C)** Population vector (PV) correlation matrix of familiar and intermediate (left) or novel (right) sessions in control (top), Δ LEC_GLU_ (middle), Δ LEC_GLU&GABA_ (bottom) (embedded figures report min:max correlation values). **(D)** Tuning curve correlation pearson coefficients between familiar and intermediate (solid line) and novel (dashed line) sessions for control (black), Δ LEC_GLU_ (green), Δ LEC_GLU&GABA_ (magenta) (control, n = 67 ROIs, paired-T test, p < 0,001; Δ LEC_GLU_, n = 97 ROIs, Wilcoxon signed-rank test, p < 0.001; Δ LEC_GLU&GABA_, n = 92 ROIs, Wilcoxon signed-rank test, p < 0.001; fam vs int: Kruskal-Wallis ANOVA, p < 0.001, Dunn-Holland-Wolfe *post hoc* test, control vs Δ LEC_GLU_, p = 0.039, control vs Δ LEC_GLU&GABA_, p = 0.021, Δ LEC_GLU_ vs Δ LEC_GLU&GABA_, p = 0.961; fam vs nov: Kruskal-Wallis ANOVA, p = 0.003, Dunn-Holland-Wolfe *post hoc* test, control vs Δ LEC_GLU_, p = 0.051, control vs Δ LEC_GLU&GABA_, p = 0.657, Δ LEC_GLU_ vs Δ LEC_GLU&GABA_, p = 0.248).

We identified spatially-tuned CA3 PN somas and dendrites in the familiar environment and compared their remapping in the subsequent environments separately in control, Δ LEC_GLU_, and Δ LEC_GLU&GABA_ (Figure 6A-B). In control conditions, CA3 PN place maps were similar for the familiar and intermediate environments, indicating pattern completion in the intermediate context (Figure 6B-D). In contrast, exposure to the novel environment caused substantial remapping of CA3 PN place maps, suggesting discrimination between the familiar and novel contexts hence pattern separation (Figure 6B-D). Silencing LEC_GLU_ decreased remapping of CA3 PN somas and dendrites, as evidenced by higher correlations of tuning curves and population vectors between familiar and intermediate (Figure 6B-D, Figure S8) as well as novel environments (Figure 6B-D, Figure S8) compared to control, whereas silencing LEC_GLU&GABA_ resulted in mixed levels of remapping, with familiar vs novel correlations similar to control (Figure 6B-D, Figure S8) but familiar vs intermediate correlations close to Δ LEC_GLU_ (Figure 6B-D, Figure S8). Altogether, these results indicate that LEC_GLU_ promotes remapping in CA3, possibly by conveying novelty-related sensory information directly to CA3 as well as recruiting orthogonalized DG inputs to CA3. Besides, LEC_GABA_ appears to increase or decrease CA3 remapping depending on whether context similarity is high or low, respectively.

These *in vivo* findings are mechanistically consistent with our *ex vivo* data showing that LEC_GABA_ promotes the integration of LEC_GLU_ with RC to yield CA3 output. However, extrapolating circuit mechanisms gathered in a reduced system to explain complex population-level dynamics would be an oversimplification. To address this, we built an attractor model of the CA3 network to explore how the changes in population dynamics resulting from our manipulations of LEC would impact pattern separation and completion *in silico*, agnostic to the intricate functional connectivity of the LEC to CA3 circuit. For this, we used the proportion of active CA3 neurons and their activity rates observed during GOL navigation in control, Δ LEC_GLU,_ and Δ LEC_GLU&GABA_ conditions (Figure 7). Irrespective of the contextual condition (familiar, intermediate, novel), we observed a general trend for the fraction of place cells to be increased with Δ LEC_GLU_ and Δ LEC_GLU&GABA_ compared to control, suggesting sparser spatial coding in control (Figure 7A-H, Figure S9, Table S2-3). These spatially tuned ROIs showed no difference in activity rates (Figure 7A-H, Figure S9, Table S2-3). Conversely, both the fraction of active non-place cells and their activity rates were decreased with Δ LEC_GLU_ compared to both control and Δ LEC_GLU&GABA_ (Figure 7A-H, Figure S9, Table S2-3), suggesting a decrease in sparsely coded contrast of CA3 spatial representations with LEC_GLU_ silencing.

**Figure 7.**
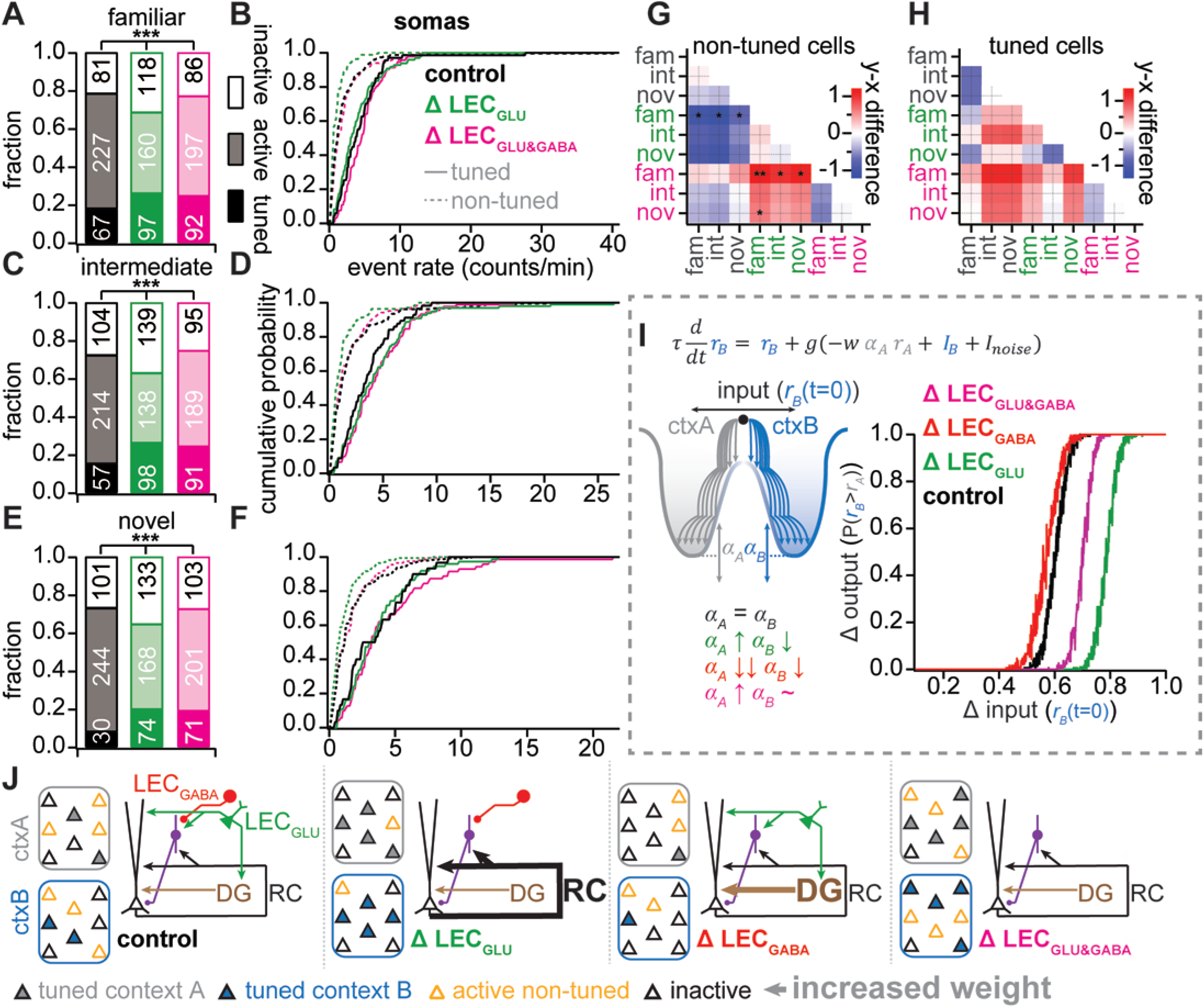
LEC_GLU_ & LEC_GABA_ bias on CA3 activity towards pattern separation & completion, respectively. **(A)** Fraction of spatially tuned ROIs, active non-spatially tuned ROIs, and inactive ROIs during the familiar session in control (black), Δ LEC_GLU_ (green), and Δ LEC_GLU&GABA_ (magenta) (fraction of tuned ROIs: χ^2^ test, p = 0.165; fraction of active non-tuned ROIs: χ^2^ test, p = 0.002). **(B)** Event rate of non-spatially tuned ROIs (dashed lines) and spatially tuned ROIs (solid lines) in the familiar session (event rate of non-tuned ROIs: control, n = 227, Δ LEC_GLU_, n = 160, Δ LEC_GLU&GABA_, n = 197, Kruskal-Wallis ANOVA, p < 0.001; event rate of tuned ROIs: control, n = 67, Δ LEC_GLU_, n = 97, Δ LEC_GLU&GABA_, n = 92, Kruskal-Wallis ANOVA, p = 0.020). **(C)** Same as (A) but for the intermediate session (fraction of tuned ROIs: χ^2^ test, p = 0.010; fraction of active non-tuned ROIs: χ^2^ test, p < 0.001). **(D)** Same as (B) but for the intermediate session (event rate of non-tuned ROIs: control, n = 214, Δ LEC_GLU_, n = 138, Δ LEC_GLU&GABA_, n = 189, Kruskal-Wallis ANOVA, p = 0.002; event rate of tuned ROIs: control, n = 57, Δ LEC_GLU_, n = 98, Δ LEC_GLU&GABA_, n = 91, Kruskal-Wallis ANOVA, p = 0.206). **(E)** Same as (A) but for the novel session (fraction of tuned ROIs: χ^2^ test, p < 0.001; fraction of active non-tuned ROIs: χ^2^ test, p = 0.007). **(F)** Same as (B) but for the novel session (event rate of non-tuned ROIs: control, n = 244, Δ LEC_GLU_, n = 168, Δ LEC_GLU&GABA_, n = 201, Kruskal-Wallis ANOVA, p < 0.001; event rate of tuned ROIs: control, n = 30, Δ LEC_GLU_, n = 74, Δ LEC_GLU&GABA_, n = 71, Kruskal-Wallis ANOVA, p = 0.598). **(G)** Difference matrix of non-spatially tuned CA3 PN somas event rates. **(H)** Difference matrix of spatially tuned CA3 PN somas event rates. **(I)** Δ output, ie neural dissimilarity (fraction of simulations yielding higher activity of the B population active in the novel context compared to the A population active in the familiar context) as a function of Δ input, ie context dissimilarity (initialization bias towards activity in the B population active in the novel context) in the attractor network model. **(J)** Schematic interpretation.

We used this information to parametrize a computational model of CA3 with two populations of spatially tuned neurons, A and B, that suppress each other and are maximally active in the familiar and novel environments, respectively. As the environment is progressively morphed from familiar (*r_B_*(t=0) = 0.1) to novel (*r_B_*(t=0) = 1.0) (Δ input), the fraction of trials with population B (*r_B_*) activity exceeding that of A (*r_A_*) (Δ output) sharply increases from 0 (stability) to 1 (discrimination) (Figure 7I). Simulating the Δ LEC_GLU_ condition by increasing the fraction of place cells and decreasing the fraction of active non-place cells resulted in increased stability (Figure 7I), hence replicating our experimental observations. Increasing the fraction of place cells alone to simulate the Δ LEC_GLU&GABA_ condition also yielded an increase of stability but to a lesser extent than with Δ LEC_GLU_ (Figure 7I). To gain further insight into the role of LEC_GABA_, we then used the model to predict what would happen with silencing of LEC_GABA_ alone (Δ LEC_GABA_), which could not be achieved under our experimental conditions. Since our *ex vivo* electrophysiology data demonstrate an overall disinhibitory action of the LEC_GABA_ input to CA3, we hypothesized that the Δ LEC_GABA_ condition would decrease both the fraction of place cells and the fraction of active non-place cells. With this, we found a decrease of stability consistent with removal of RC disinhibition by LEC_GABA_ (Figure 7I), although we note that such increased discrimination required the reduction in active place cells to exceed that of non-place cells. Altogether, these results indicate that subtle differences in CA3 PN population activity brought about by LEC_GLU_ and LEC_GABA_ can substantially bias CA3 function towards pattern separation or completion, respectively (Figure 7J).

## Discussion

### Summary of findings

Here, we investigated how entorhinal cortex long-range glutamatergic and GABAergic projections drive neuronal dynamics in area CA3 to flexibly support both pattern separation and completion. We first used freely moving behavioral assays to show that chemogenetic silencing of CA3-projecting LEC_GLU_ neurons alone or with LEC_GABA_ impaired novel object discrimination. This suggested that LEC_GLU_ and LEC_GABA_ play complementary roles in episodic learning and associated behavioral output. Next, *in vivo* two-photon calcium imaging showed that silencing LEC_GLU_ increased the proportion of active CA3 PN soma and dendrites in the population while reducing activity at the single neuron level. Further, silencing LEC_GLU_ decreased the remapping of CA3 place cells upon exposure to partially similar or completely novel environments. However, eliminating both the LEC_GLU&GABA_ together reset population activity rates and increased place cell remapping towards control states in response to the same external input manipulation. This indicated that counteracting forces may be at play in the LEC-driven modulation of CA3 activity. Therefore, we performed optogenetic circuit mapping using *ex vivo* patch-clamp electrophysiology of soma and dendrites to decipher the LEC-CA3 circuit. With dual color optogenetics, we assessed the individual and conjunctive contributions of the LEC_GLU_ and LEC_GABA_ inputs to shaping CA3 PN and IN activity. We found that LEC_GLU_ recruited a significant amount of feedforward inhibition, thus preventing CA3 PN spike output. Conversely, the LEC_GABA_ input exclusively targeted a subset of CA3 INs, thereby suppressing this LEC_GLU_-driven feedforward inhibition specifically in the somatic compartment. Hence, we found that LEC_GABA_ acts as a disinhibitory gate to selectively boost the integration of LEC_GLU_ and RC inputs to yield CA3 PN somatic output. Consistent with our *in vivo* observations, these results suggested that the LEC_GABA_-CA3 circuit interaction is poised to promote pattern completion in CA3 population dynamics. We built a computational model constrained by our population imaging and circuit mapping data to test this prediction by simulating progressively dissimilar environments. Our attractor network model replicated our experimental findings with LEC_GLU_ and LEC_GLU&GABA_ silencing and further simulated LEC_GABA_ silencing to show that a hypothesized reduction of place cells in the CA3 neuronal population would lead to decreased stability in this condition. Thus, in line with our experimental circuit-driven hypothesis, our modeling predicts that LEC_GABA_ promotes CA3 pattern completion. Taken together, our multicompartment-level experimental circuit dissection, imaging, and computational modeling reveal that long-range circuit interactions between LEC and hippocampal CA3 provide a context-dependent bidirectional switch between stability and flexibility of spatial representations through modulation of excitation, inhibition, and disinhibition of the CA3 recurrent network.

### LEC inputs impact novelty-driven behaviors

Cortico-hippocampal interactions are crucial for episodic memory and spatial navigation. While several studies have begun exploring the role of MEC in supporting hippocampal representations^68^ as well as spatial- and temporal-coding^69^, the role of neighboring area LEC still needs to be discovered. This may be partly due to the non-spatial and more abstract types of information that LEC codes for. Recordings and imaging studies in LEC of mice and humans show that, while LEC has weak spatial selectivity, this region shows context-dependent responses^30^ to a host of stimuli. These range from sensory and local cues like odors^35,37^ as well as objects or their displacement^31^, to abstract features including timing^70^ and task sequence^32^ as well as learning rules^38^, and further to behaviorally salient signals encompassing novelty^30^, punishments, rewards^36,38^ and navigational goals^71^. Few studies have directly explored the role of LEC in hippocampal memory-related behavior using a loss-of-function approach. Nevertheless, those revealed important functions for LEC inputs in odor-context associative learning^72^ and memory^73^ in CA1 and DG^74^, as well as fear context discrimination and novel object recognition in CA1^36^. Finally, LEC lesions have been found to impair the rate remapping of CA3 place cells induced by shape or color changes of the environment^42^. These findings have pivoted LEC as a central candidate for supporting novelty encoding and context-dependent remapping of neural representations in the hippocampus. Here, we showed the complementary role of glutamatergic and GABAergic LEC inputs to area CA3 in supporting object recognition by facilitating discrimination between familiarity and novelty. Indeed, lesion studies had previously shown that LEC is necessary for novel object recognition^57–59^, hence we provide further insight revealing that the LEC-CA3 component of the entorhinal-hippocampal circuit, is necessary for NOR, possibly through LEC_GLU_-conveyed object-novelty signals driving discrimination in CA3.

### Circuit operations underlying stability and flexibility of hippocampal representations

Spatial representations formed and stored within the auto-associative sub-region CA3 of the mouse hippocampus provide a rich substrate to study pattern separation and completion. A large body of experimental and theoretical work has highlighted the ability of CA3 place cells to remap in novel or morphed environments and predict how CA3 performs neuronal pattern separation and completion computations^1–6,9,10,12,13,15–18,20,49^. The local and long-range circuit interactions we decipher in our study provide a critical functional link between CA3 neural and behavioral input-output patterns.

Under control conditions during goal-oriented spatial navigation in contextually rich environments, we observed high correlations in the place ensemble across sessions when animals were re-exposed to a similar environment but a considerable amount of remapping when exposed to a novel environment. Silencing LEC_GLU_ alone resulted in more remarkable similarity or generalization, implying that LEC_GLU_ inputs promote pattern separation during transitions from familiar to novel environments in service of better discrimination. However, when LEC_GLU_ was silenced together with LEC_GABA_, the network reverted to differentiating between familiar and novel input patterns. This suggests that conjunctive LEC_GLU_ and LEC_GABA_ inputs bias the CA3 network towards pattern completion, helping generalize across similar contexts. Thus, the LEC_GLU_ and LEC_GABA_ inputs provide a dynamic range for CA3 to transition between pattern separation and completion states.

Further, by uncovering the circuit correlates involved in setting the balance between pattern separation and completion in CA3, we provide additional evidence in favor of a long-hypothesized model of CA3 function requiring both orthogonalization through the DG-CA3 path and association via EC-CA3 synapses^6^. Finally, our findings that LEC_GLU_ and LEC_GABA_ conjunctly drive CA3 activity but bidirectionally influence discrimination versus generalization of CA3 representations have implications for the interpretation of previous lesion or silencing studies (some of which were seminal to this work, see^42^) as well as for the design of future experiments. Indeed, these overlapping drives yet distinct roles of LEC_GLU_ and LEC_GABA_ highlight the importance of parsing out the molecular makeup of long-range projections to decipher inter-regional communication and associated functions in the brain.

### Deciphering CA3 *in vivo* activity dynamics through LEC input circuit mapping

In our study, combining *ex vivo* circuit dissection and *in vivo* perturbations revealed that a central feature of the LEC drive onto CA3 is the LEC_GLU_-recruited feedforward inhibition which prevented LEC_GLU_ from evoking action potential firing of CA3 PNs. LEC_GABA_ relieves CA3 PNs from this feedforward inhibition to drive recurrent network output through disinhibition. In our imaging experiments, the observed increase in the number of active CA3 PNs with LEC_GLU_ silencing (Figure 2E-I) speaks to the weight of LEC_GLU_-driven feedforward inhibition in controlling and sparsifying CA3 PN activity. This may effectively suppress background noise to generate sparsely coded contrast in CA3 ensemble activity. Thus, without the LEC_GLU_-driven feedforward inhibition, the context-dependent contrast in CA3 active ensembles is reduced (Figure 2G, 2J) which in turn may contribute to the over-generalization and decreased remapping between contexts when we silence LEC_GLU_ (Figure 6). Tied together with our behavioral findings that LEC input is important for novelty detection, perhaps the silencing of LEC_GLU_ deprives CA3 ensembles of a novelty signal about salient contextual sensory cues from the environment^30,31,33–37,42^. On the other hand, LEC_GABA_ decreases such LEC_GLU_-driven feedforward inhibition, preferentially boosting CA3 RC feedback circuit to yield somatic output. This may contribute to the decreased remapping and over-generalization observed with silencing of LEC_GLU_, pending LEC_GABA_ activity remains similar in the absence of local LEC excitatory inputs. It is possible that LEC_GABA_ projection neurons are driven by LEC_GLU_ projection neurons and hence biological removal of the latter would affect activity in both pathways. We partially addressed this with modeling of CA3 attractor network dynamics where we could isolate the hypothesized contribution of LEC_GABA_ projections and thereby reveal their role in stabilizing ensemble representations. In summary, we reveal the interplay between feedforward inhibition recruited by LEC_GLU_ and disinhibition mediated by LEC_GABA_ that provide a circuit substrate underlying the dynamics of pattern separation and pattern completion in CA3. LEC input can support high contrast CA3 place cell sparse ensembles in distinct contexts through direct excitation of a subset of CA3 PNs and feedforward inhibition of the majority of CA3 PN population. Therefore, our silencing results could explain how area CA3 place cells remap in novel environments relying on the discriminatory LEC_GLU_ signal, while maintaining stability through generalization. Thus, through an eloquent and tightly-controlled dialogue between excitation, inhibition and disinhibition, we propose that LEC_GLU_ and LEC_GABA_ act together to regulate hippocampal place cell stability and flexibility.

Further, the reduction of remapping dynamic range in CA3 PN ensembles observed with LEC_GLU&GABA_ silencing (Figure 6) may stem from an imbalance of LEC_GLU_-conveyed novelty signals leading to decreased context discrimination, and LEC_GABA_-aided pattern completion yielding similar remapping when the difference in sensory cues between environments is large (fam vs nov) but recruiting abnormally overlapping ensembles in similar environments (fam vs int). The latter observation suggests that LEC_GABA_ alone can only modulate but not override the LEC_GLU_ signal in area CA3. This is consistent with NOR performance being similarly impaired by LEC_GLU_ +/− LEC_GABA_ silencing (Figure 1D-H). It has been reported that learning a context discrimination-based fear memory task results in the emergence of a sparse and highly synchronized ensemble of neurons in area CA2/CA3, which are preferentially recruited by anterior cingulate prefrontal cortex inputs during memory retrieval^75^. Our study shows how increased signal-to-noise ratio building contrast and sparsification of active place ensembles can be mechanistically achieved through the dynamic interplay of LEC_GLU_-driven excitation and feedforward inhibition overall favoring context discrimination. Future work is needed to resolve the brain state- and task demand-dependent specific contributions of LEC_GLU_ vs LEC_GABA_ to shaping CA3 synaptic plasticity and network dynamics relevant for learning and memory recall.

### Compartment- and pathway-specific gating of activity

We found that the silencing effects on the activity of dendrites and soma *in vivo* mirror each other, perhaps an effect of backpropagated action potentials dominating our GCaMP-based calcium signals. Conversely, our *ex vivo* experiments resolved that the integrative action of LEC_GLU_ and LEC_GABA_ inputs on gating recurrent activity in CA3 is compartment specific, boosting somatic but not dendritic activity. An explanation for this can be inferred from our circuit mapping experiments where we showed that the LEC_GABA_ input targets CA3 SLM VIP+ and CCK+ neurons. It is possible these include putative VIP+/CCK+ basket cells as described in CA1^65,76,77^, thus producing perisomatic but not dendritic disinhibition. It remains to be explored whether LEC_GABA_ projections target other IN populations such as neurogliaform cells which may be relevant for dendritic disinhibition at different time scales through their volume-release of GABA causing slower shunting and hyperpolarization through GABA_B_R activation^78,79^.

Another explanation for this compartment-specific effect is that somatic spike output can occur relatively independently of distal dendritic excitation. CA3 PN dendrites show suprathreshold excitability in the form of sodium and NMDA spikes, but also backpropagated action potentials. Although we did not probe these nonlinear events in the context of synaptic plasticity, we note that CA3 NMDA spikes evoked by strong coincident activation of DG and RC are required to induce input timing-dependent plasticity^80^, and CA3 backpropagated action potentials are critical for spike timing-dependent plasticity^81^. Considering somato-dendritic propagation of nonlinear events: while dendritic NMDA spikes may facilitate somatic output^80,82^ and conversely backpropagated action potentials strongly depolarize proximal dendrites^81,83^, the compartmentalization of CA3 PNs is such that the excitability of distal dendrites is rather governed by PSP-evoked sodium dendritic spikes^83^. Hence, it is possible that our *in vivo* imaging captured mostly proximal CA3 PN dendrites where the GCaMP6f-reported calcium signal would be dominated by either backpropagated action potentials or conjunctive DG- and RC-triggered NMDA spikes, thus mirroring somatic activity. In contrast, our *ex vivo* patch-clamp recordings targeted distal CA3 PN dendrites (∼250 µm from the soma) are more likely to capture local sodium dendritic spikes evoked by integration of LEC_GLU_ and RC, rather than backpropagated action potentials or DG input which impinges on proximal dendrites.

Although LEC long-range inhibitory projections to area CA1 had been described earlier^36^, the use of dual color optogenetics allowed us to test their effect onto area CA3 together with the long-range excitatory projections from the same upstream region. This not only permits a comprehensive examination of both LEC long-range inputs but also lends region specificity and timing consistency, which is relevant to the operation of the entorhinal-hippocampal circuit since different entorhinal regions drive the hippocampus at different theta phases and gamma frequencies^60,84,85^. Further, optogenetic activation of LEC inputs enabled us to resolve their compartment-specific effects on CA3 PN somas and dendrites, which are rarely investigated^80–83^ and never in an input-selective manner. Because LEC axons impinge onto the distal dendrites of CA3 PNs, probing the consequences of LEC stimulation with somatic and dendritic recordings uncovered differential drives of each compartment through dendritic integration with local inputs and recruitment of specific IN populations. Finally, this approach also allowed us to establish rather than infer that LEC_GABA_ do not contact CA3 PN dendrites and that strong activation of LEC_GLU_ may recruit the DG feedforward and CA3 feedback local circuits. These circuit-level findings not only benefit the interpretation of our *in vivo* manipulations but also provide ground truth evidence useful for future modelling work.

### Contrasting sub-region-specific microcircuitry and related input-output transformations

Our findings both reveal unifying principles and draw contrast with the functional organization of entorhinal long-range GABAergic projections to area CA1. Firstly, similar to LEC^36^ and MEC^43^ GABAergic inputs to CA1, we find that LEC_GABA_ exert an overall disinhibitory action on area CA3. We further provide evidence that LEC_GABA_ exclusively target CA3 INs but not PN distal dendrites, which is inferred but not resolved in area CA1. As for key differences, the LEC_GABA_ input to CA1 gates dendritic spikes in contrast to the LEC_GABA_ input to CA3 gating somatic spikes. This is due to the LEC_GABA_ innervation of CA1 SR/SLM CCK+ dendrite-targeting INs that control dendritic excitability and associated synaptic plasticity without affecting somatic output in CA1 PNs^36^. Whether molecular differences between CA1 and CA3 INs and/or dendritic domains specifically route the LEC_GABA_ path towards dendritic versus somatic disinhibition warrants further investigation. Alternatively, long-range inhibitory projections may differentially affect non-recurrent circuits such as CA1 and recurrent excitatory networks such as CA3 which are commonly found in neocortical areas. Regardless, we note that cortical long-range GABAergic projections targeting of local INs seems to be conserved across brain regions^44,45,47^, although the net effect is not systematically disinhibitory^46^.

Interestingly, this preferential and often reciprocal long-range GABAergic innervation of local INs may help synchronizing distant brain areas^44^. This is because local inhibitory transmission is particularly effective at pacing network oscillations^86–89^, thereby amplifying the effect of long-range GABAergic transmission across regions. This has notably been demonstrated in the neocortical networks as well as the septo-hippocampal and MEC-CA1 circuits across several oscillation patterns^43,46,90–95^. Although currently unexplored, this may apply to LEC as well since LEC_GABA_ targets theta-modulated INs in the form CCK+ INs in CA1 and putative VIP+ / CCK+ INs in CA3. Another surprising and perhaps common feature of long-range GABAergic cells is their high propensity to respond to salient sensory stimuli regardless of the modality^36,96^. This may serve the encoding of coincident multimodal inputs and thereby contribute to novelty detection. Indeed, silencing of either the general LEC_GABA_ hippocampal projections^36^ or the LEC_GLU&GABA_ to CA3 input impair novel object recognition, which might reflect a default in encoding novelty such that familiar objects are not registered. Further roles for long-range GABAergic transmission in learning and memory include encoding of object location^46,93^, spatial learning^97^, and contextual memory^97–99^.

### Specific contributions of LEC input to CA3 pattern separation and completion

Our findings are consistent with and build on the long recognized role of CA3 in balancing pattern separation and completion functions^1–6,9,10,12,13,15–18,20,49^. The recurrent connectivity architecture of the CA3 network lends itself to attractor dynamics which stabilize activity of neuronal ensembles representing a given point in the feature space by sharing mutual excitation while distributing inhibition onto the rest of the population^20^. Such network organization produces invariant output when presented with similar inputs since the activation of subparts of the ensemble is sufficient to activate the remainder of the ensemble, hence performing pattern completion^100^. This is thought to rely on reactivation of previously formed CA3 PN ensembles by EC-driven sensory information^6^. On the other hand, larger differences in input will force activity to shift towards another stable point in the attractor network, thereby yielding pattern separation. This is would be brought about LEC-driven novelty signals since LEC neurons readily exhibit decorrelated activity upon sensory information mismatch^5^, as well as the sparsified DG input which may randomly select subsets of CA3 PNs to represent a given context^20^. Our attractor network model encompasses both ends of such behavior, allowing us to recapitulate our experimental observations, provide mechanistic interpretations, and extrapolate further to non-experimentally-tested conditions.

Our head-fixed goal-oriented learning task combined with *in vivo* two-photon CA3 population imaging allowed us to probe remapping in gradually dissimilar sensory contexts as a proxy for pattern separation and completion. Compared to our baseline remapping in control, silencing of LEC_GLU_ produced decreased remapping thus suggesting increased pattern completion. This was accompanied with an increase of place cells (without changes of place cell activity rate) and a decrease of active non-place cells (which also exhibited reduced activity rates). Implementing these changes of population dynamics in our attractor network model reproduced our experimental findings. Given our understanding of the LEC-CA3 circuit, we interpret these results as a consequence of decreased LEC_GLU_-driven feedforward inhibition being permissive for increased place cell expression, which in turn suppresses non-place cell activity through feedback inhibition locally in CA3. With an increased fraction of cells contributing to the “familiar” attractor and lower synaptic noise in the network (reduced non-place cell activity), it follows that stability of activity in the “familiar” basin would increase. Conversely, we hypothesize from our *ex vivo* experiments that silencing of LEC_GABA_ should reduce disinhibition as well as the weight of recurrent excitatory connections in CA3. This would result in increased local inhibition and decreased recurrent excitation, overall leading to lower non-place cell activity and reduced place cell propensity. Our model shows that, as long as the latter exceeds the former, the reduced strength and span of attractors in the CA3 population facilitates transitions from stable points in the feature space, thus promoting pattern separation. Finally, with an increase in place cell propensity (ie more neurons contributing to any given attractor) together with unchanged non-place cell activity (ie a synaptic noise similar to control) observed in the LEC_GLU&GABA_ silencing condition, our model produces a modest increase in pattern completion. This is consistent with decreased CA3 remapping upon small but not large context changes in the condition. We infer that silencing of both LEC_GLU_ and LEC_GABA_ reduces feedforward inhibition hence facilitating place cell expression, while also dampening the weight of recurrent connectivity thus preventing excessive feedback inhibition and thereby striking an offset balance between increased attractor stability (via increased neuronal contribution) and retained tonic non-place cell activity (via unchanged synaptic noise). Altogether, our combined *ex vivo* circuit dissection, *in vivo* population imaging, and *in silico* attractor dynamics allows to formulate a parsimonious yet comprehensive mechanistic explanation for the complex influence of LEC onto CA3 pattern separation and completion functions.

### Resource availability

#### Lead contact

Further information and requests for resources and reagents should be directed to and will be fulfilled by the lead contact, Jayeeta Basu.

#### Materials availability

This study used the AAV2/7:h56D-ChR2-sfGFP custom-designed viral reagent, courtesy of Boris Zemelman. Please contact the corresponding author for further details.

#### Data and code availability

All data reported in this paper will be shared by the lead contact upon request. All original code that has been deposited on github is available to reviewers and will be publicly available as of the date of publication. Any additional information required to reanalyze the data reported in this paper is available from the lead contact upon request.

### Experimental model and study participants details

#### Mice

All experiments were conducted in accordance with the National Institute of Health guidelines and with the approval of the New York University School of Medicine Institutional Animal Care and Use Committee (IACUC). Mice were obtained from Jackson Laboratory and subsequent breeding was established in-house. Acute slice electrophysiology experiments used GAD2-Cre, PV-Cre, SST-Cre, CCK-Cre, VIP-Cre, Ai9 and Ai14 transgenic mice from both sexes, 8 weeks-to 6 months-old, with a C57BL/6J genetic background. *In vivo* 2-photon calcium imaging experiments used GAD2-Cre x Thy1-GCaMP6f (GP5.5^101^) transgenic mice from both sexes, 3 to 6 months-old, with a C57BL/6J genetic background. Freely moving behavioral experiments used C57BL/6J mice from both sexes, 3 to 12 months-old.

### Method details

#### Virus preparation

An adeno-associated virus encoding an inhibitory neuron promoter h56D^102^, channelrhodopsin2(H134R)-sfGFP, woodchuck hepatitis virus posttranscriptional regulatory element (WPRE) and SV40 polyadenylation signal was assembled using a modified helper-free system (Stratagene) as a serotype 2/7 (rep/cap genes) AAV and harvested and purified over sequential cesium chloride gradients as previously described^103^. The codon-optimized channerhodopsin2 fusion protein included an EAGAVSGGVY linker between the protein domains and C-terminal Golgi and endoplasmic reticulum export signals^104,105^ to aid membrane expression.

#### Stereotaxic surgery

Animals (4+ weeks-old) were anaesthetized with isofluorane (1.5-3 %, inhaled) and buprenorphine (0.1 mg/kg, injected intra-peritoneally). Mice were placed in a stereotaxic apparatus (Stoelting), a small incision (∼0.5 cm length) was made in the skin to expose the skull, and the skull was levelled flat according to bregma and lambda. A small craniotomy (∼0.5 mm diameter) was drilled in the skull above the injection sites and 276-414 nL of virus was injected (Drummond Scientific Nanoject II) into the brain at the following coordinates: anterior–posterior relative to bregma: 3.2 and 3.0 mm for LEC, 1.9 mm for CA3; medial-lateral relative to midline: 4.5 and 4.7 mm for LEC, 2.2 mm for CA3; dorsal-ventral relative to the surface of the brain: 2.5, 2.8 and 3.0 mm for LEC, 1.8, 2.0 and 2.2 mm for CA3. The injection pipette was slowly lowered into the brain up to 0.2 mm deeper than the deepest targeted coordinate, left at this location for 1 min, retracted to the actual coordinate and held there for 30 s prior to injection. 46-69 nL of virus was injected in 23 nL increments spaced by 15 s at each z coordinate, with an additional 2 min pause between z coordinates and a final 10 min incubation at the last (shallowest) z coordinate of each injection site before slowly retracting the injection pipette out of the brain. Mice were sutured, given neosporin topically on the incision site, injected with 1mL of sterile saline subcutaneously, monitored until full recovery from anesthesia was observed, and given analgesic post-operative care (buprenorphine, 0.1 mg/kg, injected intra-peritoneally) for 3 days. Infection sites were confirmed to be specific *post hoc* for all experiments. The adeno-associated virus (AAV) AAV2/5:EF1α-DIO-hChR2(H134R)-eYFP-WPRE-HGHpA (Addgene #20298) was used at 1×10^13^ vg/mL, AAV2/5:CaMKIIa-hChR2(H134R)-eYFP (Addgene #26969) was used at 1.5×10^13^ vg/mL, AAV2/5:CaMKII-Chronos-GFP (Neurophotonics #319) was used at 1.3×10^13^ vg/mL, AAV2/5:Syn-flex-ChrimsonR-tdTomato (Addgene #62723) was used at 4.74×10^12^-2×10^13^ vg/mL, AAV2/7:h56D-ChR2-sfGFP (kindly provided by Boris Zemelman) was used at 2×10^13^ vg/mL, AAV2/5:hSyn-DIO-hM4D(Gi)-mCherry (Addgene #44362) was used at 1.4×10^13^ vg/mL, AAV2/5:CaMKII-flex-hM4D(Gi)-mCitrine (Addgene #50477) was used at 1.1×10^13^ vg/mL, AAV2/5:CaMKII-PSAM4-GlyR-IRES-eGFP (Addgene #119744) was used at 2.5×10^13^ vg/mL, AAV2/5:Syn-DIO-eGFP (Addgene #50457) was used at 7×10^12^ vg/mL, AAV2/9:Dlx-flex-TdTomato (Addgene #83894) was used at 7×10^12^ vg/mL, AAVretro:Syn-Cre (Neurophotonics #1299) was used at 1.4×10^13^ vg/mL.

#### Electrophysiology

Artificial cerebrospinal fluid (ACSF) and protective dissection ACSF (dACSF) were oxygenated with a 95 % O_2_ and 5 % CO_2_ mixture at all times. Mice were deeply anaesthetized with isoflurane (5 % for 5 min, inhaled) and perfused transcardially with ∼20 mL of ice-cold NMDG-based dACSF^106^ containing (in mM): NMDG 93, KCl 2.5, NaH_2_PO_4_ 1.25, NaHCO_3_ 30, HEPES 20, glucose 25, thiourea 2, Na-ascorbate 5, Na-pyruvate 3, CaCl_2_ 0.5, MgCl_2_ 10. Brains were then rapidly removed, cortico-hippocampal complexes were dissected out and placed upright into a custom-made agar mold in ice-cold dACSF. 400 µm thick transverse slices were cut (vibratome, Leica VT1200S) at low speed (0.04 mm/s) and blade vibration amplitude (0.5 mm) in ice-cold dACSF. Slices were transferred to an immersed-type holding chamber and maintained in ACSF containing the following (in mM): NaCl 125, KCl 2.5, NaH_2_PO_4_ 1.25, NaHCO_3_ 25, glucose 22.5, Na-ascorbate 1, Na-pyruvate 3, CaCl_2_ 2, MgCl_2_ 1. Slices were incubated at 32 °C for ∼20 minutes and then maintained at room temperature for at least 30 minutes prior to recordings. Individual slices were transferred to a recording chamber perfused with ACSF at 3-5 mL/min (peristaltic pump, Watson Marlow) at 30 °C (in-line heater, Warner Instruments TC-324B). Tissue was visualized under an upright microscope (Zeiss Examiner A1 or Olympus BX51WI) equipped with DIC or Dodt gradient contrast at 5x-40x magnification with additional zoom optics 1-2.5x, and captured by a video camera (Hamamatsu ORCA-spark or ORCA-flash4.0). The headstages connected to the recording electrodes were mounted on motorized micromanipulators (Luigs & Neumann GmbH). Patch-clamp recordings were performed with potassium- or cesium-based intracellular solution containing the following (in mM): K- or Cs-methyl sulfonate 135, KCl 5, EGTA-KOH 0.1, HEPES 10, NaCl 2, MgATP 5, Na2GTP 0.4, Na2-phosphocreatine 10, and either Alexa 594 (50-100 µM) or biocytin (4 mg/mL). GΩ seal were formed and whole-cell recordings were obtained from CA3 pyramidal neurons soma (blind patch) or dendrites (blind patch) and interneurons (visual patch). Somatic patch pipette resistances were 2-5 MΩ, series resistances were 8-20 MΩ. Dendritic patch pipette resistances were 13-17 MΩ, series resistances were 40-60 MΩ. Series resistance was compensated 0-50 % to amount to an 8-10 MΩ actual resistance in voltage-clamp (somatic recordings). Bridge-balance was applied in current-clamp. Unless stated otherwise, the membrane potential was held at −70 mV in current-clamp. The liquid junction potential was < 10 mV and not corrected for. ChR2-, Chronos- or ChrimsonR-expressing LEC inputs were stimulated optically with 1-10 ms-long 470 or 625 nm light pulses of intensity 0-100 % (0.0-4.47 mW/mm^2^). Dual-color optogenetic experiments were performed with Chronos- and ChrimsonR-expressed in LEC_GLU_ and LEC_GABA_, respectively stimulated with 1 ms 1-2 % (0.07-0.14 mW/mm^2^) 470 nm and 10 ms 100 % (4.47 mW/mm^2^) 625 nm light pulses that were empirically determined to allow spectral separation (intensity below cross-talk threshold of ChrimsonR activation by 470 nm light). Monosynaptic transmission from ChR2- or Chronos-expressing LEC_GLU_ was probed with 1ms 1-2 % (0.1-0.14 mW/mm^2^) 470 nm light pulses that were empirically determined to elicit monophasic and TTX & 4-AP resistant EPSCs in CA3 PNs, as well as below population spike initiation threshold from LFP recordings in DG GC and CA3 SP layers. Conversely, poly-synaptic transmission from ChR2- or Chronos-expressing LEC_GLU_ was probed with 100% (3.76 mW/mm^2^) 470 nm light pulses that were empirically determined to elicit polyphasic and TTX & 4-AP sensitive EPSCs in CA3 PNs, as well as above population spike initiation threshold from LFP recordings in DG GC and CA3 SP layers. The medial CA3 recurrent input and proximal dentate gyrus mossy fiber input were stimulated electrically with ACSF-filled pipettes mounted on manual micromanipulators (Siskiyou) and placed in CA1 *stratum radiatum* (SR) to antidromatically activate CA3 and in the hilus near the border of the upper blade granule cell layer to directly activate DG axons, respectively. Post-synaptic responses were evoked with constant current stimulation units (Digitimer Ltd.) delivering 0.1 ms long 25-200 µA current pulses through monopolar electrodes. Pharmacological agents were added to ACSF at the following concentrations (in µM): 10 NBQX and 50 D-APV to block AMPA and NMDA receptors, 1-2 SR95531 and 2 CGP55845A to block GABAA and GABAB receptors, 10 clozapine N-oxide (CNO) to activate hM4D(Gi) DREADDs, 0.2-1 tetrodotoxin (TTX) to prevent sodic action potential generation, 100 4-aminopyridine (4-AP) to block KV1 potassium channels, 1-10 (2S,2’R,3’R)-2-(2’,3’-dicarboxycyclopropyl)-glycine (DCG-IV) to inhibit glutamate release from mossy fiber terminals by activating mGluR2/3. Data was obtained using a Multiclamp 700B amplifier (Molecular Devices), sampled at 10 kHz, digitized using a Digidata 1550B AD/DA board (Molecular Devices), and acquired with the pClamp 10 software (Molecular Devices). Data analysis was performed in IgorPro (Wavemetrics) with custom-written code.

#### Two-photon calcium imaging

All anesthesia, analgesia and pre-surgical procedures were performed as described above (see stereotaxic injections). GAD2-Cre x Thy1-GCaMP6f expressing the GCaMP6f calcium fluorescent indicator in glutamatergic neurons were previously injected unilaterally with AAV2/5:hSyn-DIO-hM4D(Gi)-mCherry and AAV2/5:CaMKII-PSAM4-GlyR-IRES-eGFP in LEC. 3 weeks later, a circular 3 mm diameter craniotomy was made centered on the ipsilateral CA3 imaging coordinates (anterior–posterior relative to bregma: 1.4 mm; medial-lateral relative to midline: 1.6 mm). The skull fragment was removed, and a vacuum system was used to gently aspirate the overlying cortex and external capsule. Ice-cold ACSF was used to irrigate the area throughout the duration of the procedure. A cranial window (3 mm diameter, 1.7 mm length stainless steel cannula attached to 3 mm diameter glass coverslip) was then implanted over the area. The window was sealed to the skull using Vetbond, and a custom-designed 3D-printed plastic headpost was cemented over the skull. Mice were allowed to recover for 3-5 days before being placed under water restriction after which their weight was monitored daily to ensure it remained at least 80 % of baseline. Mice were head-fixed under the two-photon microscope on a treadmill belt and trained to run for 5 % sucrose water as described previously^66,67^. *In vivo* two-photon imaging was performed using a dual galvanometric and resonant laser scanning two-photon microscope (Ultima, Bruker), coupled to a tunable Ti:Sapphire laser (MaiTai eHP DeepSee, Spectraphysics) pulsed at a 80 MHz repetition rate and < 70 fs pulse width along with dispersion compensation. GCaMP6f fluorophore was excited at 920 nm, using a resonant scanning X-galvanometer (8 kHz) paired with a 6 mm standard scanning Y-galvanometer. The scanning system was mounted on a movable objective Ultima microscope, equipped with an orbital nosepiece coupled to a 16 X, 0.8 NA, 3 mm water immersion objective (Nikon) and a piezo drive for angled imaging and ultrafast volumetric scanning. Imaging was performed at a scan speed of 29 fps, using 512 x 512 frame size (1.085 µm/pixel resolution). Fluorescence signal was detected using high-sensitivity GaAsP photomultiplier tubes (model 7422PA-40 PMTs, Hamamatsu). CA3 PN calcium transients reported by GCaMP6f were imaged on a single stable field of view in each mouse throughout the experiments. Mice were first imaged for a single 10 min-long session daily as they ran on the same textured belt (familiar) for water rewards delivered at random positions on the belt (random foraging, RF) for 9 days. On days 10, 11 and 12, mice were subjected to a 10 min RF imaging session on the familiar belt and allowed to rest for 1 h before another 10 min session on the familiar belt immediately followed by a final 10 min session on a novel belt (a new belt was used for each day). Mice were injected with either saline, PSEM817 (0.3 mg/kg, ip), or PSEM817 (0.3 mg/kg, ip) and CNO (5 mg/kg, ip) during the 1 h rest period between the 2 familiar sessions of days 10, 11 and 12. Mice were then trained for 2 days (days 13 and 14) to learn a fixed reward location at 90-100 cm (goal-oriented learning, GOL) on the familiar belt with the addition of a tone (4 kHz) and odor pulse (10 % pentyl acetate in mineral oil) interleaved and delivered at 0.25 Hz throughout the session. On days 15, 16 and 17, mice were injected with saline, PSEM817, or PSEM817 and CNO 30 min before being subjected to a 40 min imaging session encompassing a sequence of 10 min familiar (familiar), 10 min intermediate, 10 min familiar (familiar 2), and 10 min novel as described next. The familiar sessions consisted of running on the familiar belt with the reward location at 90-100 cm and the same tone (4 kHz) and odor (10 % pentyl acetate in mineral oil) as before. The intermediate session was run on the familiar best but with reward location at 150-160 cm and a different tone (10 kHz) and odor (10 % (+)-α-pinene in mineral oil). The novel session used a novel belt (a new belt was used for each day) with reward location at 30-40 cm and the unfamiliar tone (10 kHz) and odor (10 % (+)-α-pinene in mineral oil). Reward location was baited by manual water delivery in the vicinity of the reward zone if mice failed to find the reward location after 5 min within each session to encourage running every 2 laps until mice licked to receive water. Environmental stimuli delivery and behavioral data recording were performed with custom-designed Arduino hardware. Imaging data was acquired with the PrairieView software (Bruker) and synchronized with the Arduino inputs and outputs. For each mouse, image sequences from all sessions were concatenated and processed by suite2p^107^ for motion correction and ROI detection. ROI sets were thereafter manually curated and annotated using ImageJ^108^. All subsequent analyses were performed in MATLAB (MathWorks) with custom-written code. For each ROI (spatial component), the activity trace (time component) was obtained by averaging across all pixels within the ROI at each time point (frame) and normalized to baseline fluorescence. Calcium transients were detected and demixed from overlapping ROIs using the fitness algorithm of the d-NMF package^109^. Briefly, activity traces were detrended and z-scored to detect putative events as positive peaks overshooting a threshold of 2 standard deviations from baseline. Putative events were then screened using a correlation threshold of 0.27 between the spatial component of the ROI and the actual spatial footprint of the ROI in the movie frame at the time of the event. Subsequent analysis was performed on these curated events, with further behavioral restriction to run epochs as defined by a speed of the animal > 2 cm/s. Spatially-resolved activity rates were computed by smoothing and dividing the event counts by the occupancy within 5 cm-wide bins. For each session, spatial information content^110^ was computed for each ROI as:

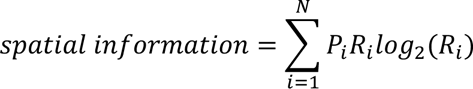

where *P_i_* is the normalized occupancy in the *i*^th^ spatial bin such that the sum across all *P_i_* = 1, and *R_i_* is the normalized value of the activity rate in the *i*^th^ spatial bin such that the sum across all *R_i_* = 1. Statistical significance of spatial tuning was determined for each ROI within each session by shifting event times from −250 to 250 time bins and normalizing the spatial information content at zero shift to that of the 99^th^ percentile at non-zero shift. ROIs meeting criteria of a normalized spatial information content > 1 and a number of events > 4 were considered as significantly spatially tuned. Spatial tuning curves were parametrized by circularly smoothing rate maps with a sigma of 15 cm and fitting the resulting smoothed curve with multiple Von Mises functions increasing in number until the residuals fell below 25 % of the maximal value of the original rate map.

#### Computational modelling

We simulated a mean-field model of two population *r_A_* and *r_B_*. They followed the given dynamics

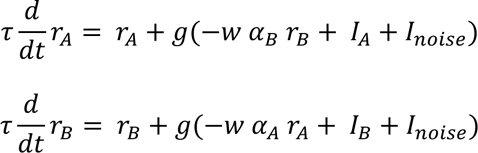

where *τ* = 10 [a.u.] is the time constant of the dynamics, *I_A_* = 1.05 and *I_B_* = 0.95 are the currents intro the *r_A_* and *r_B_* population, respectively and *I_noise_* is a noise drawn from a uniform distribution between −0.2 and 0.2. *w* = 1 is the cross-population inhibitory weight. The function *g*(*x*) = 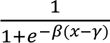 is a sigmoid with parameters *β* = 10 and *γ* = 0.5. Due to the fact that the *r_A_* receives a larger current, it is the active population in the familiar environment. To simulate the effect of the environment becoming progressively dissimilar, we initialized *r_B_* to different values from 0.1 to 1.0 (Δ input, in increments of 0.001) and at the same time *r_A_* = 1 − *r_B_*. We simulated the network for 100 time steps [a.u.] and for 100 trials. We reported the percentage of times *r_B_* > *r_A_* at the end of the simulation as a measure of neural difference (Δ output). To simulate the different manipulations, we set *α_A_* = *α_B_* = 1 for the control condition. For the Δ LEC_GLU_ condition which experimentally (i) increased the fraction of place cells and (ii) decreased the fraction of active non-place cells (Figure 7, Supplemental Figure 9), we increased the modulation factor *α_A_* = 1.3 and decrease the factor *α_B_* = 0.9. For the Δ LEC_GLU&GABA_ condition which increased the fraction of place cells (Figure 7, Supplemental Figure 9), we only increased the modulation factor *α_A_* = 1.3. For the Δ LEC_GABA_ condition, we assumed that (i) the fraction of active non-place cells would decrease, therefore reducing the factor *α_B_* = 0.9 and (ii) and that the fraction of place cells would decrease more, thus further reducing *α_A_* = 0.8. Code is available to reviewers (github).

#### Freely moving behavior

C57BL/6J mice previously injected bilaterally with AAVretro:Syn-Cre into CA3 and either AAV2/5:Syn-DIO-eGFP, AAV2/5:CaMKII-flex-hM4D(Gi)-mCitrine or AAV2/5:hSyn-DIO-hM4D(Gi)-mCherry into LEC were allowed to recover for 3 weeks before being handled (including scruffing) for 5 min daily by the experimenter for 1 week prior to behavioral testing. All experiments were conducted in presence of background white noise (70 dB) and white light (150-300 lux). The open field apparatus consisted of an opaque white rectangular arena (length 37 cm, width 29 cm, height 27 cm). Mice were first habituated to the empty open field arena for 10 min for 2 days. The novel object recognition (NOR) task comprised of a first session where mice were injected with CNO (5 mg/kg, ip) 30 min before being exposed to the open field arena with 2 identical objects placed ∼7cm away from the walls on each side of the rectangular area for 10 min (encoding), before being returned to their home cage for 1 h. Mice were then exposed to the open field arena with one object identical to the encoding session and the other one replaced by a new object of similar size, both at their original locations, for 10 min (recall). The Barnes maze consisted of an opaque black circular platform (90 cm diameter) elevated 70 cm from the floor with 20 holes (5 cm diameter) located 40 cm from the center and equidistant along the perimeter, with a single hole leading to a hidden escape box. The maze was surrounded locally (10 cm from platform) by a circular opaque white curtain containing 5 equidistant movable local cues on ∼75 % of its perimeter, and distally (100 cm from platform) by an opaque black curtain providing a fixed distal cue spanning ∼25 % of its perimeter. Training trials consisted of mice being placed in the escape box for 1 min, then moved into an opaque black cylinder (12.5 cm diameter) in the center of the maze for 10-15 s before the cylinder was removed and mice were let free to explore the maze until they found the hole containing the escape box (goal) for a maximal duration of 3 min (encoding) after which mice were gently guided to the goal if unable to find it until then. Test trials were similar to training trials except mice were placed directly into the center of the maze inside the opaque black cylinder at the beginning of the trial and were not guided to the goal afterwards (recall). The experiment consisted of 4 repetitions of encoding trials spaced by 15 min with CNO injection 30 min prior to the 1st trial for 6 days (days 1-6), and a single drug-free recall trial on the 7th day (day 7). All trials were recorded with an infrared camera (Basler, 5 acA640-100gm) placed atop the open field arena or Barnes maze platform. Video data was sampled at 50 Hz and acquired with the Pylon software (Basler). For each batch of mice and each behavioral paradigm (NOR, Barnes maze), videos were visually examined, trimmed and cropped if necessary. Curated videos were then concatenated and processed with DeepLabCut^111^ to track the XY position of the head (marked at the intersection of the lateral line between the ears and the rostro-caudal line between the nose and the neck) and body (marked at the intersection of the lateral line between the front paws and the rostro-caudal line between the neck and the tail protrusion from the body). Tracking data was analyzed with custom-written code in IgorPro (Wavemetrics) to yield the relevant metrics. We computed the mouse’s position, distance travelled, speed, head-body angle, and travel direction across time. From these metrics, we parsed the data into behaviorally relevant indices: total trial length, epochs inside target subdivisions of the area (2 halves of the open field for NOR, 4 quadrants centered around the goal escape hole for Barnes maze), epochs inside vs outside of target zones (12.5 cm diameter around objects for NOR, 8 cm diameter around escape holes for Barnes maze), epochs inside target zones with the animal facing the target irrespective of travel direction (Figure S2B-C, +/− 45° angle towards objects for NOR, +/− 45° angle towards goal escape hole for Barnes maze), epochs leading to the first visit of target zones, number of visits to each target zone, and epochs in the center vs periphery of the area (33×26 cm inner rectangle of the open field for NOR, 72 cm diameter inner circle for Barnes maze; as a control for thigmotaxis). The discrimination index was computed as:

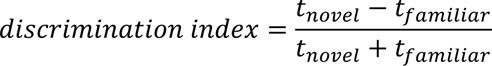

where *t_novel_* and *t_familiar_* are the times spent interacting with the novel and familiar objects, respectively.

#### Histology

Mice were deeply anaesthetized with isoflurane (5 % for 5 min, inhaled) and perfused transcardially with ∼20 mL of PBS followed by ∼20 mL of 4 % PFA in PBS. Brains were removed and fixed in 4 % PFA in PBS at 4 °C for at least 24 h. Brains were washed in 0.3 M Glycine in PBS for 15 min followed by 3 x 15 min washes in PBS. 100 µm thick coronal slices were cut (vibratome, Leica VT1000S) and stored in PBS at 4 °C. Similarly, samples recovered from *ex vivo* electrophysiology (400 µm thick transverse slices) were fixed in 4 % PFA in PBS at 4 °C for at least 24 h, washed in 0.3 M Glycine in PBS for 15 min followed by 3 x 15 min washes in PBS, and stored in PBS at 4 °C. Tissue was permeabilized with 0.5 % Triton in PBS (PBST) for 2 x 20 min, blocked with 10 % NGS in 0.5 % PBST for 4 h, and incubated with primary antibodies in 3 % NGS 0.1 % PBST overnight at 4 °C. Tissue was then washed with 0.2 % PBST for 15 min followed by 3 x 30 min washes in PBS before being incubated with secondary antibodies and Streptavidin where applicable in 3 % NGS 0.1 % PBST for 24 h (100 µm slices) or 48 h (400 µm slices) at 4 °C. Lastly, slices were washed 5 x 15 min in PBS and mounted in Vectashield Hard Set Mounting Medium with DAPI (Vector Laboratories). Primary antibodies were rabbit anti-GFP (1:1000, ThermoFisher #A6455), chicken anti-GFP (1:1000, Abcam #13970). All secondary antibodies and dyes were purchased from ThermoFisher: Alexa Fluor 488-conjugated goat anti-rabbit (1:1000, #A11008), Alexa Fluor 488-conjugated donkey anti-chicken (1:1000, #A78948), Alexa Fluor 488-conjugated goat anti-chicken (1:1000, #A11039), Alexa Fluor 555-conjugated streptavidin (1:500, #S11227), Alexa Fluor 594-conjugated streptavidin (1:500, #S21381), Alexa Fluor 647-conjugated streptavidin (1:500, #S21374), Alexa Fluor 455-conjugated Neurotrace (1:200, #N21479). Samples were screened by epifluorescence imaging (Olympus VS200). Relevant samples were imaged with an inverted Zeiss Axio Observer Z1 confocal microscope using 10x (air, 0.3 NA), 20x (air, 0.8 NA) or 40x (oil, 1.3 NA) objectives (Zeiss) and 405, 488, 594, and 647 nm lasers for fluorophore excitation. Images were acquired as 1024 x 1024 pixels 16 bits Z-stacks with a 5 µm (10x, 20x) or 1 µm (40x) Z-step size and tiled in X and Y as needed to cover the samples, and thereafter stitched using the Zen Microscopy software (Zeiss). Further processing of confocal or epifluorescence images was done in ImageJ^108^.

### Quantification and statistical analysis

#### Statistics

Results are reported ± SEM. Normality was tested with the Shapiro-Wilk and Jarque–Bera tests, and homoscedasticity was tested using Barlett’s test to choose between parametric and nonparametric statistical analysis. Statistical significance was assessed using the χ^2^ test, Student’s T test, Mann-Whitney U test, paired Student’s T test, Wilcoxon signed-rank test, one-way ANOVA, Kruskal-Wallis ANOVA, repeated-measures ANOVA, Friedman ANOVA, Tukey *post hoc* test, Nemenyi *post hoc* test, Dunn-Holland-Wolfe *post hoc* test, or two-way ANOVA where appropriate. The symbols *, **, and *** denote P values <0.05, <0.01, and <0.001, respectively.

### Contributions

JB and VR conceived the project and designed experiments; VR performed all the experiments and analysis; KO assisted with the stereotaxic injections, *ex vivo* patch-clamp recordings, and immunohistochemistry related to the long-range GABAergic projection to CA3 interneuron circuit mapping experiments; RGD and CDJ performed the freely moving behavioral testing and related immunohistochemistry; SKR performed the CA3 cranial window imaging preparation and assisted with the *in vivo* two-photon imaging data acquisition; BVZ provided the AAV2/7:h56D-ChR2-sfGFP construct; CC performed computational modeling; VR and JB wrote the paper with input from all authors; JB acquired funding, provided resources, supervised personnel and managed the project.

## Acknowledgments

This work was supported by an NIH NINDS BRAIN INITIATIVE 1R01NS109994 grant to JB and CC, an NIH NINDS 1R01NS109362-01, an NINDS 1RM1NS132981-01, a McKnight Scholar Award in Neuroscience, a McKnight Endowment Fund for Neuroscience Mathew Pecot URM Award, a Klingenstein-Simons Fellowship Award in Neuroscience, Alzheimer’s Association Research Grant to Promote Diversity – New to the Field (AARG-D-NTF), an Alfred P. Sloan Research Fellowship, Mathers Charitable Foundation Award, a Whitehall Research Grant, an American Epilepsy Society Junior Investigator Award, a Blas Frangione Young Investigator Research Grant, a New York University Whitehead Fellowship for Junior Faculty in Biomedical and Biological Sciences and a Leon Levy Foundation Award to JB. VR was supported by a Young Researchers Bettencourt Prize, KO was supported by an NIH 5T32MH019524-30 training award, SKR was supported by an NIH T32GM007308 training grant, CJ was supported by an NIH NIMH Diversity Supplement training award 3R01MH122391-04S1 (Parent award R01MH122391PIs Buzsáki /Basu) and BVZ was supported by funding from an NIH NINDS 1U01 NS099720 and 1U01 NS094330. NYU High Performance Computing resources were used for data analysis. NYU Genotyping Core Laboratory was used for genotyping mice. We thank Martial Dufour, Roland Zemla and Jason Moore for the early development of the two-photon imaging preparation, head-fixed behavioral setup, and imaging data processing and analysis software at the Basu lab. We Melissa Hernandez-Frausto for the establishment of the freely moving behavior setups at the Basu lab, and technical advice on the behavioral paradigms. We thank Michael Long for electrophysiology equipment resources for the project and feedback on the manuscript, We are grateful to Olesia Bilash, Tanvi Butola, György Buzsáki, Melanie Druart, Simon Chamberlain, Maya Hopkins, Dayu Lin, Rebecca Piskorowski, and Richard Tsien for their input at various stages of the project and helpful comments on the manuscript.

## SUPPLEMENTAL INFORMATION

**Figure S1, related to Figure 1.**
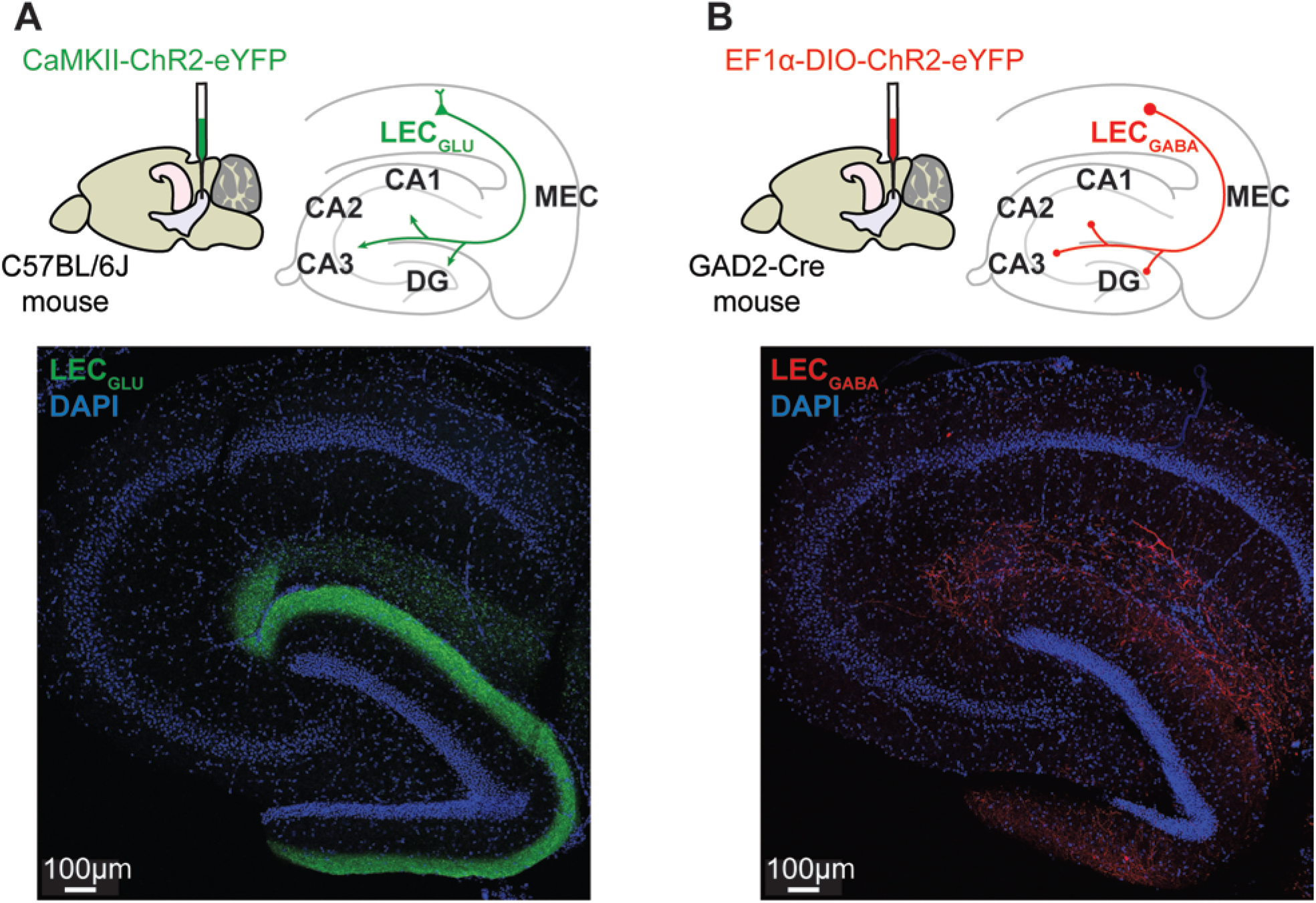
Labelling of LEC_GLU_ and LEC_GABA_ projections to CA3. **(A)** Top, viral strategy. Bottom, sample LEC_GLU_ projections (eYFP, green) to the hippocampus, with DAPI staining (blue). **(B)** Left, viral strategy. Right, sample LEC_GABA_ projections (eYFP, red) to the hippocampus, with DAPI staining (blue).

**Figure S2, related to Figure 1.**
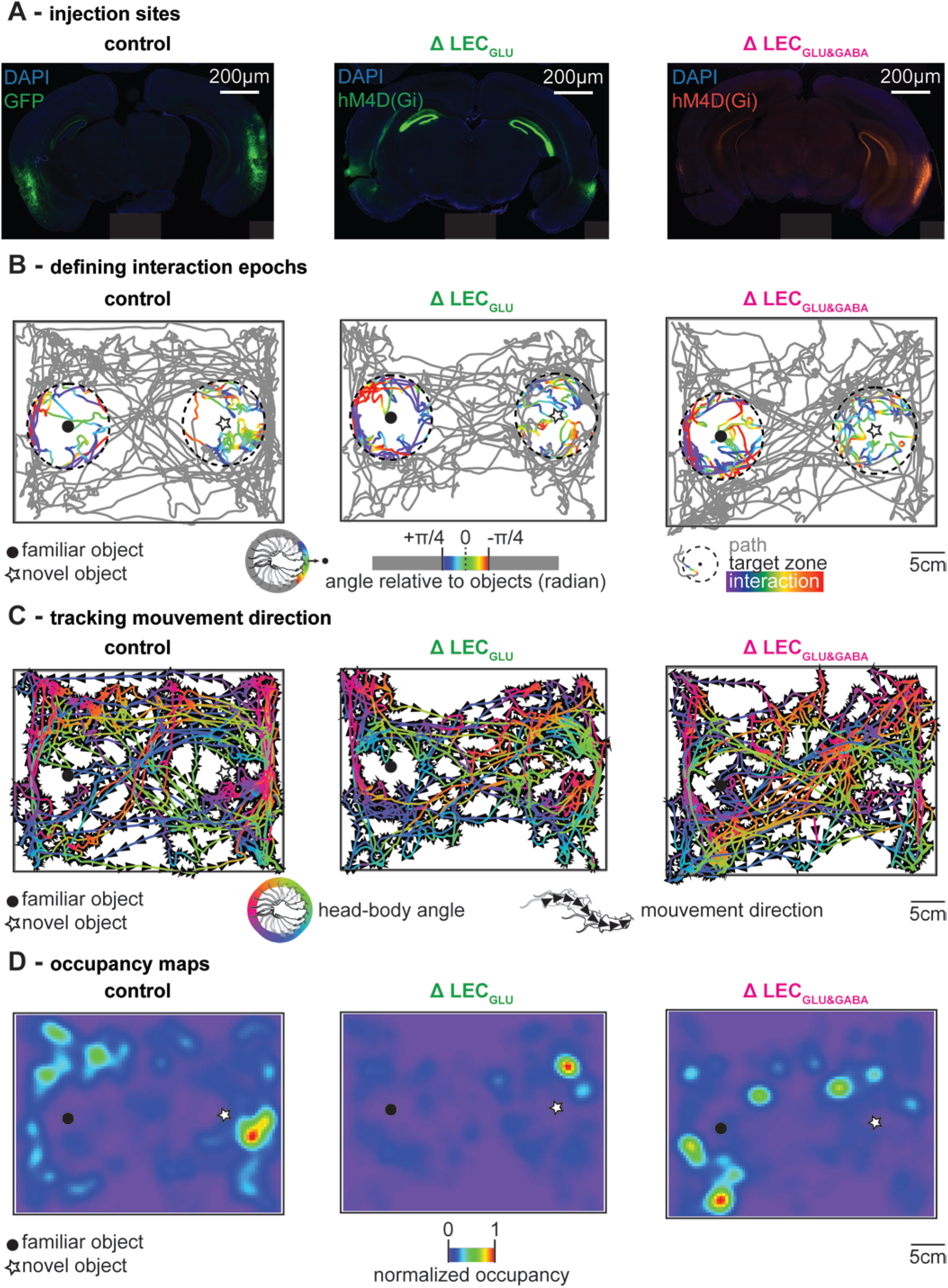
Additional NOR-related information. **(A)** Sample injection sites of AAVs targeting CA3-projecting LEC neurons, showing GFP expression in LEC_GLU_ and LEC_GABA_ neurons (GFP, green, left), hM4D(Gi) expression in LEC_GLU_ neurons alone (mCitrine, green, middle), and hM4D(Gi) expression in LEC_GLU_ and LEC_GABA_ neurons (mCherry, red, right). **(B)** Sample NOR recall sessions showing mouse path (grey) in the open field (black outline) and exploration of the familiar (closed circle) and novel (star) objects. Bouts of interaction with the objects are color-coded according to the mouse head-body angle spanning +/− 45° relative to the corresponding object ([−45°: +45°] shown in rainbow, [−180°: −45°]∪[+45°: +180°] shown in grey). **(C)** Same as (B) but with colors showing the absolute head-body angle and arrows showing the direction of travel. **(D)** Same as (B) but showing normalized occupancy maps.

**Figure S3, related to Figure 1.**
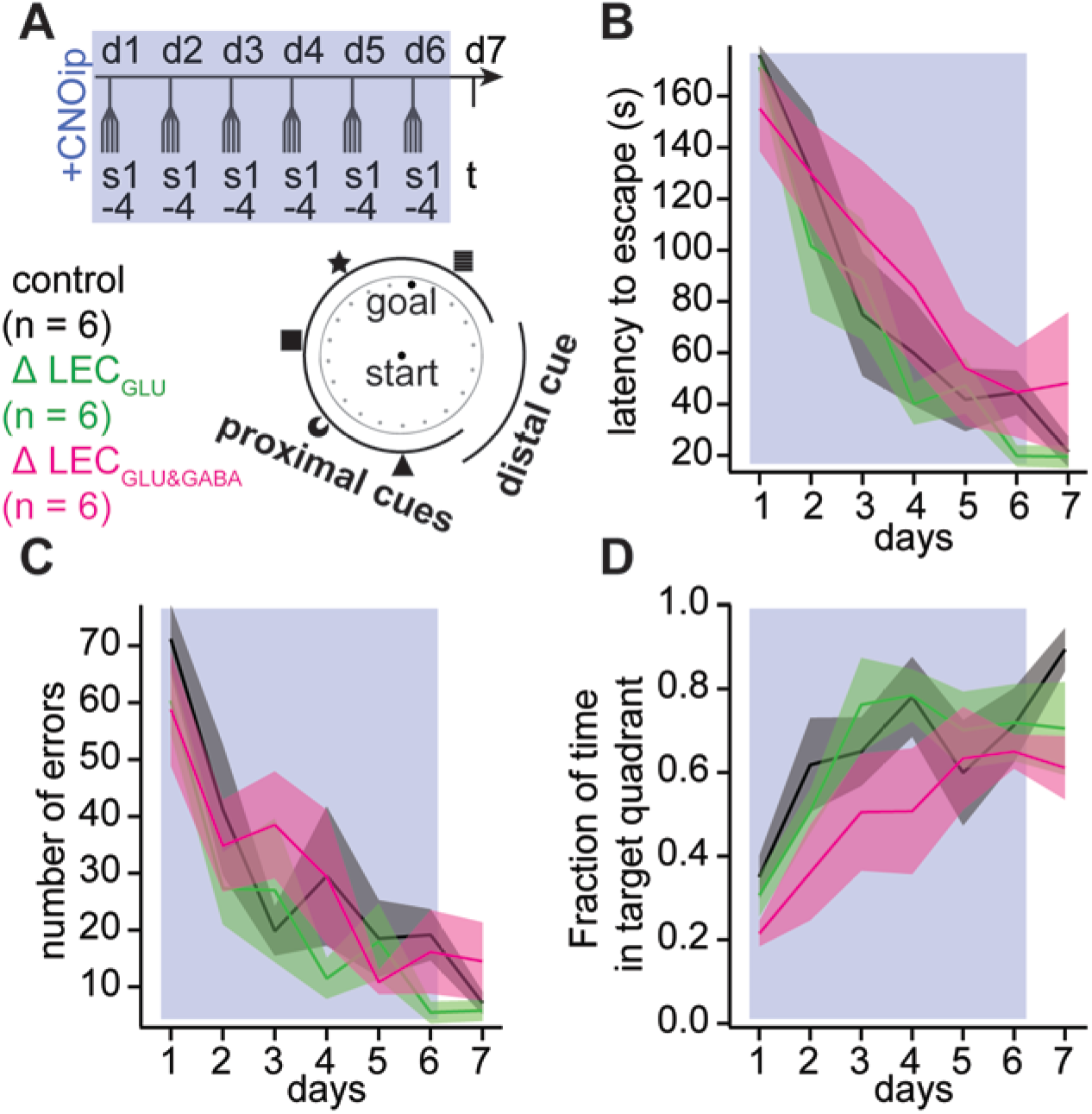
LEC silencing does not affect Barnes maze performance. **(A)** Experimental design. **(B)** Time-course of latency to escape (n = 6 each for control, Δ LEC_GLU_ and Δ LEC_GLU&GABA_, daily sessions 1-6, two-way ANOVA, treatment, p = 0.272, session, p < 0.001, treatment x session, p = 0.882; test session day 7, 21.5 ± 5.9 s for control, n = 6, 19.5 ± 4.4 s for Δ LEC_GLU_, n = 6, 48.2 ± 27.6 s for Δ LEC_GLU&GABA_, n = 6, Kruskal-Wallis ANOVA, p = 0.923). **(C)** Time-course of number of errors (n = 6 each for control, Δ LEC_GLU_ and Δ LEC_GLU&GABA_, daily sessions 1-6, two-way ANOVA, treatment, p = 0.174, session, p < 0.001, treatment x session, p = 0.657; test session day 7, 7.2 ± 2.0 for control, n = 6, 5.8 ± 1.8 for Δ LEC_GLU_, n = 6, 14.5 ± 6.9 for Δ LEC_GLU&GABA_, n = 6, Kruskal-Wallis ANOVA, p = 0.874). **(D)** Time-course of fraction of time spent in the target quadrant (n = 6 each for control, Δ LEC_GLU_ and Δ LEC_GLU&GABA_, daily sessions 1-6, two-way ANOVA, treatment, p = 0.012, session, p < 0.001, treatment x session, p = 0.859; test session day 7, 0.89 ± 0.05 for control, n = 6, 0.70 ± 0.11 for Δ LEC_GLU_, n = 6, 0.61 ± 0.08 for Δ LEC_GLU&GABA_, n = 6, one-way ANOVA, p = 0.080). Error bars represent SEM.

**Figure S4, related to Figure 3.**
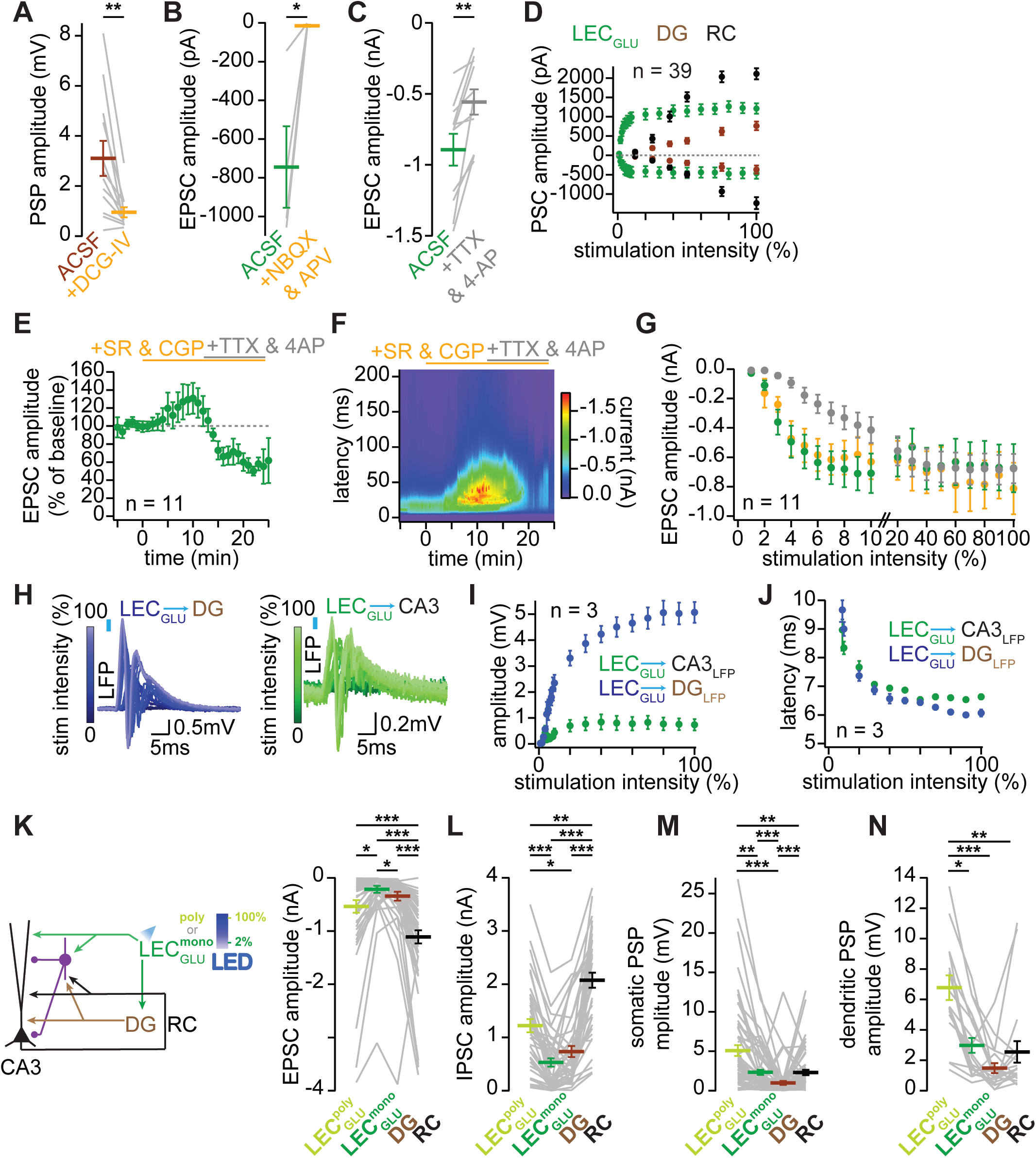
Pharmacology of synaptic transmission of interest in CA3. **(A)** DG-evoked CA3 PN PSP amplitude before and after application of DCG-IV (3.1 ± 0.7 mV in ACSF and 1.0 ± 0.2 mV in DCG-IV, n = 12, paired-T test, p = 0.003). **(B)** LEC_GLU_-evoked CA3 PN EPSC amplitude before and after application of NBQX & APV (−744.9 ± 210.0 pA in ACSF and - 14.7 ± 5.5 pA in NBQX & APV, n = 4, paired-T test, p = 0.038). **(C)** LEC_GLU_-evoked CA3 PN EPSC amplitude before and after application of TTX & 4-AP (−893.7 ± 111.8 pA in ACSF and −557.6 ± 89.5 pA in TTX & 4-AP, n = 11, paired-T test, p = 0.001). **(D)** Input-output curves of LEC_GLU_-(green, same data as Figure 3C), DG-(brown), and RC-evoked (black) CA3 PN EPSC (negative) and IPSC (positive) amplitudes. **(E)** Time-course of LEC_GLU_-evoked CA3 PN EPSC amplitude. **(F)** Time x latency x current time-course of LEC_GLU_-evoked CA3 PN EPSC. **(G)** Input-output curves of LEC_GLU_-evoked CA3 PN EPSC amplitude in ACSF (green), SR & CGP (orange), and TTX & 4-AP (grey). **(H)** Sample traces of LEC_GLU_-evoked LFP recordings in and DG GL (left, blue) CA3 SP (right, green). **(I)** Input-output curves of LEC_GLU_-evoked population spike amplitude in CA3 SP (green) and DG GL (blue), (n = 3). **(J)** Input-output curves of LEC_GLU_-evoked population spike latency in CA3 SP (green) and DG GL (blue) (n = 3). **(K)** LEC_GLU_ polysynaptic-(green), LEC_GLU_ monosynaptic-(teal), DG-(brown), and RC-evoked (black) CA3 PN EPSC amplitudes (−536.3 ±119.0 pA for LEC_GLU_ polysynaptic, −217.0 ± 65.5 pA for LEC_GLU_ monosynaptic, −343.4 ± 84.1 pA for DG, −1108.6 ± 123.5 pA for RC, n = 50, Friedman ANOVA, p < 0.001, Nemenyi *post hoc* test, LEC_GLU_ polysynaptic vs LEC_GLU_ monosynaptic, p = 0.019, LEC_GLU_ polysynaptic vs DG, p = 0.990, LEC_GLU_ polysynaptic vs RC, p < 0.001, LEC_GLU_ monosynaptic vs DG, p = 0.044, LEC_GLU_ monosynaptic vs RC, p < 0.001, DG vs RC, p < 0.001). **(L)** Same as (K) but with CA3 PN IPSC amplitudes (1221.1 ± 125.6 pA for LEC_GLU_ polysynaptic, 529.8 ± 79.6 pA for LEC_GLU_ monosynaptic, 733.2 ± 104.3 pA for DG, 2075.1 ± 141.1 pA for RC, n = 41, Friedman ANOVA, p < 0.001, Nemenyi *post hoc* test, LEC_GLU_ polysynaptic vs LEC_GLU_ monosynaptic, p < 0.001, LEC_GLU_ polysynaptic vs DG, p = 0.045, LEC_GLU_ polysynaptic vs RC, p = 0.005, LEC_GLU_ monosynaptic vs DG, p = 0.591, LEC_GLU_ monosynaptic vs RC, p < 0.001, DG vs RC, p < 0.001). **(M)** Same as (K) but with CA3 PN somatic PSP amplitudes (5.1 ± 0.7 mV for LEC_GLU_ polysynaptic, 2.3 ± 0.3 mV for LEC_GLU_ monosynaptic, 1.0 ± 0.2 mV for DG, 2.3 ± 0.3 mV for RC, n = 61, Friedman ANOVA, p < 0.001, Nemenyi *post hoc* test, LEC_GLU_ polysynaptic vs LEC_GLU_ monosynaptic, p = 0.002, LEC_GLU_ polysynaptic vs DG, p < 0.001, LEC_GLU_ polysynaptic vs RC, p = 0.003, LEC_GLU_ monosynaptic vs DG, p < 0.001, LEC_GLU_ monosynaptic vs RC, p = 0.998, DG vs RC, p < 0.001). **(N)** Same as (K) but with CA3 PN dendritic PSP amplitudes (6.8 ± 0.8 mV for LEC_GLU_ polysynaptic, 3.0 ± 0.5 mV for LEC_GLU_ monosynaptic, 1.5 ± 0.3 mV for DG, 2.6 ± 0.7 mV for RC, n = 19, Friedman ANOVA, p < 0.001, Nemenyi *post hoc* test, LEC_GLU_ polysynaptic vs LEC_GLU_ monosynaptic, p = 0.030, LEC_GLU_ polysynaptic vs DG, p < 0.001, LEC_GLU_ polysynaptic vs RC, p = 0.001, LEC_GLU_ monosynaptic vs DG, p = 0.101, LEC_GLU_ monosynaptic vs RC, p = 0.685, DG vs RC, p = 0.617). Error bars represent SEM.

**Figure S5, related to Figure 3.**
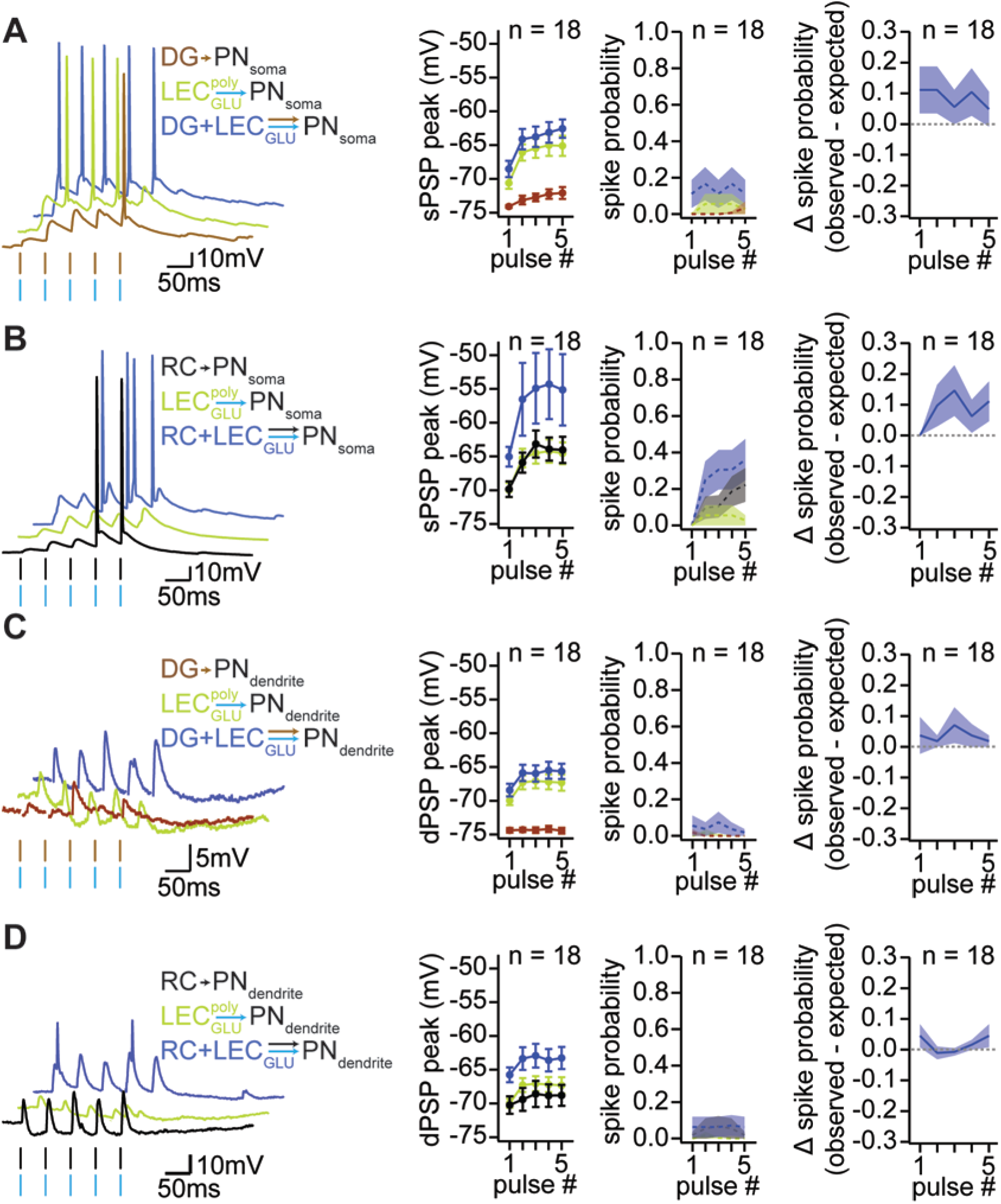
LEC glutamatergic inputs can recruit DG and RC inputs to yield CA3 PN output. **(A)** Left, sample traces of CA3 PN somatic responses to repeated stimulations of LEC_GLU_ (green), DG (brown), and LEC_GLU_ + DG (blue) inputs at 20Hz. Middle, LEC_GLU_-(green), DG-(brown), and LEC_GLU_ + DG-evoked (blue) CA3 PN somatic PSP peak (closed circles) and spike probability (dashed lines). Right, delta spike probability between observed DG + LEC_GLU_ stimulation versus expected from linear sum of these inputs (observed vs expected spike probability: n = 18, two-way ANOVA, observed vs expected, p = 0.050, pulse #, p = 0.915, observed vs expected x pulse #, p = 0.981). **(B)** Same as (A) but with RC stimulation (black) instead of DG (observed vs expected spike probability: n = 18, two-way ANOVA, observed vs expected, p = 0.146, pulse #, p = 0.008, observed vs expected x pulse #, p = 0.944). **(C)** Same as (A) but with CA3 PN dendritic responses (observed vs expected spike probability: n = 18, two-way ANOVA, observed vs expected, p = 0.074, pulse #, p = 0.871, observed vs expected x pulse #, p = 0.928). **(D)** Same as (A) but with CA3 PN dendritic responses (observed vs expected spike probability: n = 18, two-way ANOVA, observed vs expected, p = 0.599, pulse #, p = 0.967, observed vs expected x pulse #, p = 0.971). Error bars represent SEM.

**Figure S6, related to Figure 4.**
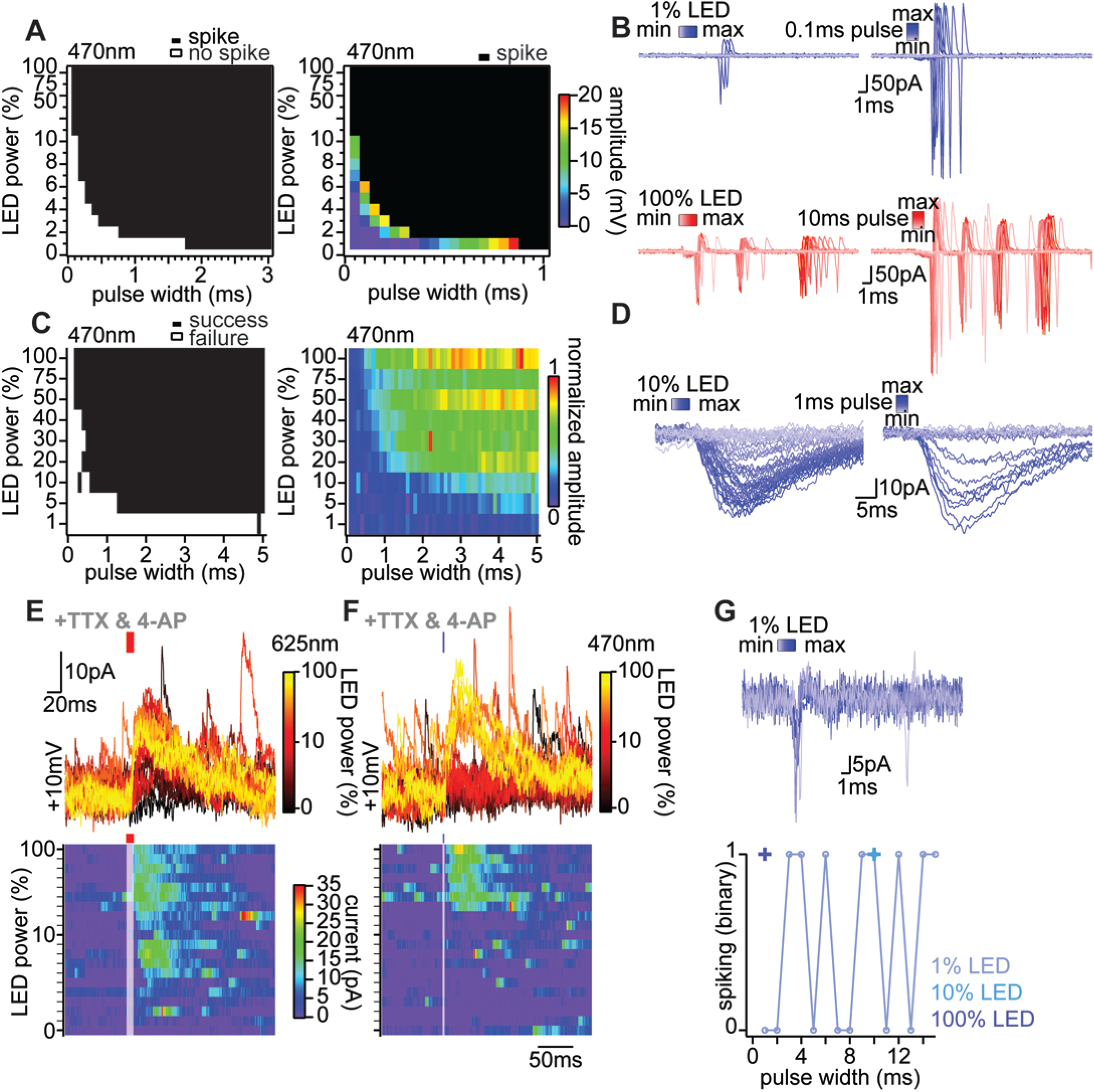
Chronos & ChrimsonR validation. **(A)** Spike probability of ChrimsonR-expressing LEC_GABA_ neurons as a function of 470 nm light pulse width vs intensity, recorded in cell-attached (left) and whole-cell (right) configuration. **(B)** Sample traces of 470 nm (top) and 625 nm (bottom) light-evoked action potential firing recorded in cell-attached configuration in ChrimsonR-expressing LEC_GABA_ neurons with increasing pulse width (left) and intensity (right). **(C)** Chronos-expressing LEC_GLU_-evoked EPSC success probability (left) and amplitude (right) recorded in CA3 PNs. **(D)** Sample traces of Chronos-expressing LEC_GLU_-evoked EPSCs recorded in CA3 PNs with increasing 470 nm light pulse width (left) and intensity (right). **(E)** Sample traces (top) and time x power x current input-output (bottom) of LEC_GABA_-evoked CA3 IN IPSCs in TTX & 4-AP, using 625 nm light to activate ChrimsonR-expressing LEC GABAergic axons. **(F)** Same as (E) but with 470 nm light. **(G)** Sample traces (top) and spiking quantification (bottom) of Chronos-expressing LEC_GLU_ neurons recorded in cell-attached configuration with increasing pulse width.

**Figure S7, related to Figure 4.**
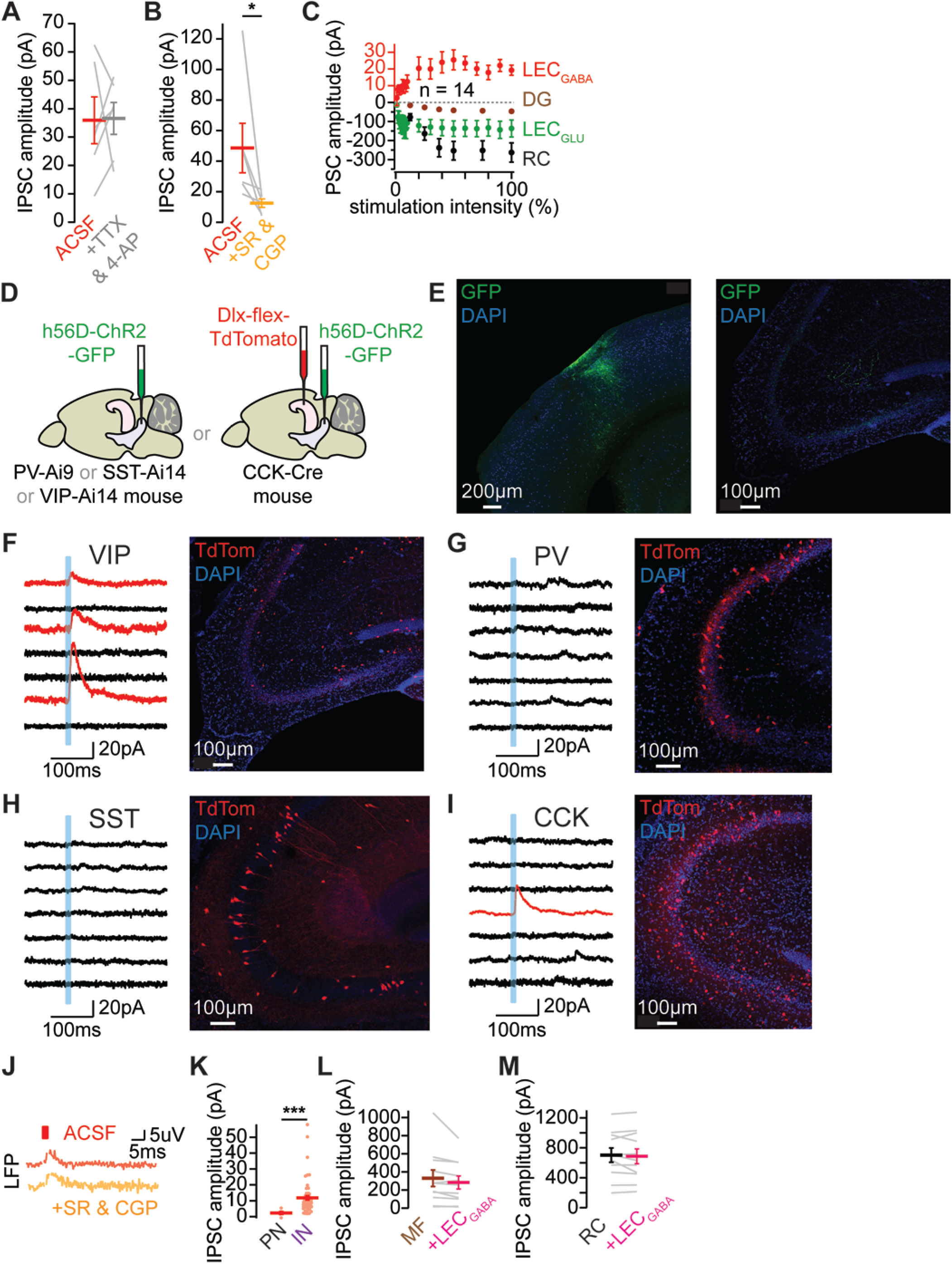
Mapping LEC_GABA_ inputs to CA3. **(A)** LEC_GABA_-evoked CA3 IN IPSC amplitude before and after application of TTX & 4-AP (35.9 ± 8.2 pA in ACSF and 36.6 ± 5.6 pA in TTX & 4-AP, n = 6, paired-T test, p = 0.945). **(B)** LEC_GABA_-evoked CA3 IN IPSC amplitude before and after application of SR & CGP (48.6 ± 16.2 pA in ACSF and 12.5 ± 2.7 pA in SR & CGP, n = 6, Wilcoxon signed-rank test, p = 0.031). **(C)** Input-output curves of LEC_GLU_-(green), LEC_GABA_-(red), DG-(brown), and RC-evoked (black) CA3 PN EPSC (negative) and IPSC (positive) amplitudes (EPSC amplitude: −391.4 ± 243.0 pA for LEC_GLU_, n = 6, −47.5 ± 8.0 for DG, n = 50, −329.8 ± 82.9 for RC, n = 39, Kruskall-Wallis ANOVA, p < 0.001, Dunn-Holland-Wolfe *post hoc* test, LEC_GLU_ vs DG, p = 0.167, LEC_GLU_ vs RC, p = 0.973, DG vs RC, p = 0.012). **(D)** Viral strategy. **(E)** Sample injection site of AAV targeting LEC_GABA_ (GFP, green, left) neurons and their projections to CA3 (right), with DAPI staining (blue). **(F)** Left, sample traces of seven CA3 VIP-expressing INs under voltage-clamp at +10mV upon LEC_GABA_ stimulation showing unresponsive (black) and responsive (red) post-synaptic cells (each trace is the average response of a single cell). Right, CA3 VIP-expressing INs (TdTomato, red) with DAPI staining (blue). **(G)** Same as (F) but with PV-expressing INs. **(H)** Same as (F) but with SST-expressing INs. **(I)** Same as (F) but with CCK-expressing INs. **(J)** Sample traces of CA3 SLM LFP upon LEC_GABA_ stimulation in ACSF (red) and SR & CGP (orange), note that the positive voltage deflection seen in ACSF remain in SR & CGP therefore likely indicating a photoelectric artifact rather than a synaptic event. **(K)** LEC_GABA_-evoked IPSC amplitude in CA3 PNs and INs (11.8 ± 1.4 pA for INs, n = 55; 2.3 ± 1.0 pA for PNs, n = 5, Mann-Whitney U test, p < 0.001). **(L)** CA3 PN IPSC amplitude evoked by DG with or without LEC_GABA_ stimulation (329.4 ± 90.9 pA for DG and 283.0 ± 71.5 pA for DG + LEC_GABA_, n = 11, paired-T test, p = 0.092). **(M)** Same as (L) but with RC (702.2 ± 95.3 pA for RC and 686.3 ± 99.6 pA for RC + LEC_GABA_, n = 11, paired-T test, p = 0.464). Error bars represent SEM.

**Figure S8, related to Figure 6.**
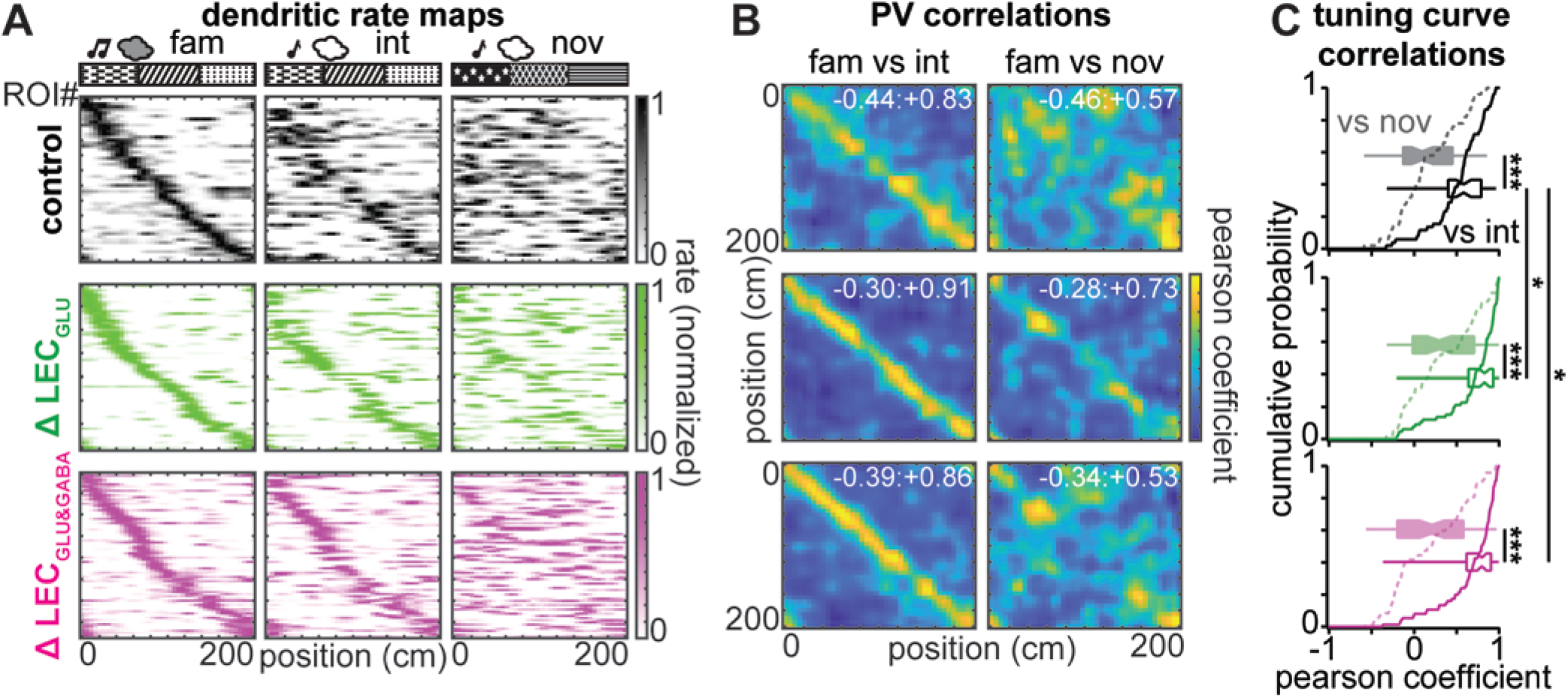
Effect of LEC_GLU_ +/−LEC_GABA_ silencing on CA3 dendritic remapping. **(A)** Normalized rate maps of CA3 place cell dendrites sorted according to their place field location in the familiar session. Left to right: familiar, intermediate, and novel sessions. Top to bottom: control (black), Δ LEC_GLU_ (green), Δ LEC_GLU&GABA_ (magenta). Topmost schematics represent experimental design. **(B)** Population vector (PV) correlation matrix of familiar and intermediate (left) or novel (right) sessions in control (top), Δ LEC_GLU_ (middle), Δ LEC_GLU&GABA_ (bottom) (embedded figures report min:max correlation values). **(C)** Tuning curve correlation pearson coefficients between familiar and intermediate (solid line) and novel (dashed line) sessions for control (black), Δ LEC_GLU_ (green), Δ LEC_GLU&GABA_ (magenta) (control, n = 55 ROIs, paired-T test, p < 0,001; Δ LEC_GLU_, n = 71 ROIs, paired-T test, p < 0,001; Δ LEC_GLU&GABA_, n = 77 ROIs, Wilcoxon signed-rank test, p < 0.001; fam vs int: Kruskal-Wallis ANOVA, p < 0.001, Dunn-Holland-Wolfe *post hoc* test, control vs Δ LEC_GLU_, p = 0.019, control vs Δ LEC_GLU&GABA_, p = 0.047, Δ LEC_GLU_ vs Δ LEC_GLU&GABA_, p = 0.894; fam vs nov: Kruskal-Wallis ANOVA, p = 0.040, Dunn-Holland-Wolfe *post hoc* test, control vs Δ LEC_GLU_, p = 0.275, control vs Δ LEC_GLU&GABA_, p = 0.991, Δ LEC_GLU_ vs Δ LEC_GLU&GABA_, p = 0.269).

**Figure S9, related to Figure 7.**
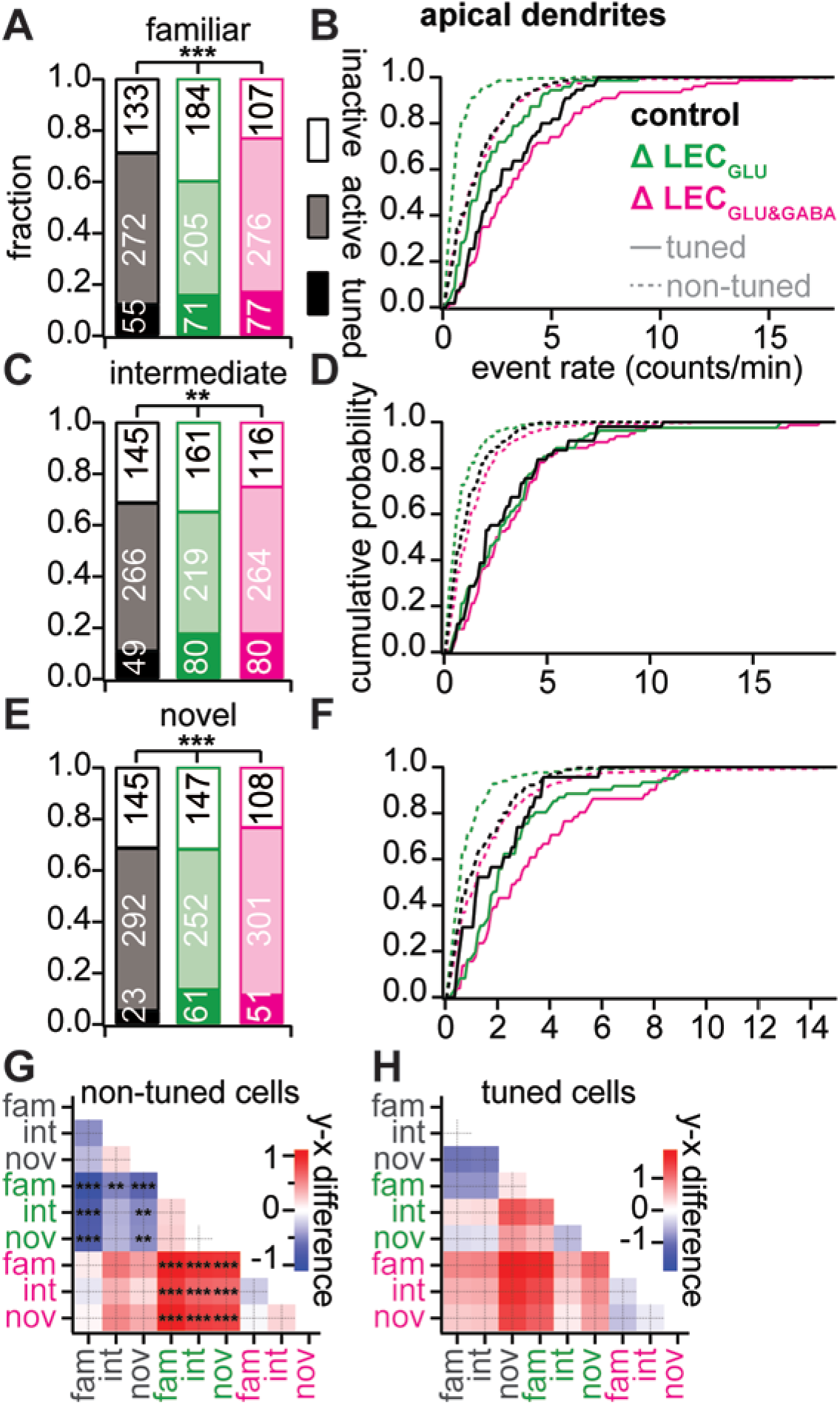
Effect of LEC_GLU_ +/− LEC_GABA_ silencing on CA3 dendritic activity during GOL. **(A)** Fraction of spatially tuned ROIs, active non-spatially tuned ROIs, and inactive ROIs during the familiar session in control (black), Δ LEC_GLU_ (green), and Δ LEC_GLU&GABA_ (magenta) (fraction of tuned ROIs: 0.12 in control, 0.15 in Δ LEC_GLU_, 0.17 in Δ LEC_GLU&GABA_, χ^2^ test, p = 0.482; fraction of active non-tuned ROIs: 0.67 in control, 0.53 in Δ LEC_GLU_, 0.72 in Δ LEC_GLU&GABA_, χ^2^ test, p < 0.001). **(B)** Event rate of non-spatially tuned ROIs (dashed lines) and spatially tuned ROIs (solid lines) in the familiar session (event rate of non-tuned ROIs: 1.6 ± 0.1 events/min in control, n = 272, 0.6 ± 0.1 events/min in Δ LEC_GLU_, n = 205, 1.6 ± 0.1 events/min in Δ LEC_GLU&GABA_, n = 276, Kruskal-Wallis ANOVA, p < 0.001; event rate of tuned ROIs: 2.9 ± 0.3 events/min in control, n = 55, 2.2 ± 0.2 events/min in Δ LEC_GLU_, n = 71, 3.8 ± 0.4 events/min in Δ LEC_GLU&GABA_, n = 77, Kruskal-Wallis ANOVA, p < 0.001). **(C)** Same as (A) but for the intermediate session (fraction of tuned ROIs: 0.11 in control, 0.17 in Δ LEC_GLU_, 0.17 in Δ LEC_GLU&GABA_, χ^2^ test, p = 0.055; fraction of active non-tuned ROIs: 0.65 in control, 0.58 in Δ LEC_GLU_, 0.69 in Δ LEC_GLU&GABA_, χ^2^ test, p = 0.039). **(D)** Same as (B) but for the intermediate session (event rate of non-tuned ROIs: 1.2 ± 0.1 events/min in control, n = 266, 0.8 ± 0.1 events/min in Δ LEC_GLU_, n = 219, 1.5 ± 0.1 events/min in Δ LEC_GLU&GABA_, n = 264, Kruskal-Wallis ANOVA, p < 0.001; event rate of tuned ROIs: 2.9 ± 0.3 events/min in control, n = 49, 3.2 ± 0.3 events/min in Δ LEC_GLU_, n = 80, 3.5 ± 0.3 events/min in Δ LEC_GLU&GABA_, n = 80, Kruskal-Wallis ANOVA, p = 0.476). **(E)** Same as (A) but for the novel session (fraction of tuned ROIs: 0.05 in control, 0.13 in Δ LEC_GLU_, 0.11 in Δ LEC_GLU&GABA_, χ^2^ test, p = 0.002; fraction of active non-tuned ROIs: 0.67 in control, 0.63 in Δ LEC_GLU_, 0.74 in Δ LEC_GLU&GABA_, χ^2^ test, p = 0.064). **(F)** Same as (B) but for the novel session (event rate of non-tuned ROIs: 1.3 ± 0.1 events/min in control, n = 292, 0.9 ± 0.1 events/min in Δ LEC_GLU_, n = 252, 1.7 ± 0.1 events/min in Δ LEC_GLU&GABA_, n = 301, Kruskal-Wallis ANOVA, p < 0.001; event rate of tuned ROIs: 1.9 ± 0.3 events/min in control, n = 23, 2.7 ± 0.3 events/min in Δ LEC_GLU_, n = 61, 3.3 ± 0.3 events/min in Δ LEC_GLU&GABA_, n = 51, Kruskal-Wallis ANOVA, p = 0.044). **(G)** Difference matrix of non-spatially tuned CA3 PN dendrites event rates. **(H)** Difference matrix of spatially tuned CA3 PN dendrites event rates.

**Table S1, related to Figure 2.** RF event rate p-values (Dunn-Holland-Wolfe *post hoc* test)

| somas | | control | | $\Delta\text{LEC}_{\text{GLU}}$ | | $\Delta\text{LEC}_{\text{GLU\&GABA}}$ | |
| --- | --- | --- | --- | --- | --- | --- | --- |
|  |  | fam | nov | fam | nov | fam | nov |
| control | fam |  |  |  |  |  |  |
|  | nov | 0.999 |  |  |  |  |  |
| $\Delta\text{LEC}_{\text{GLU}}$ | fam | 0.161 | 0.058 | | | | |
|  | nov | 0.089 | 0.028 | >0.999 |  |  |  |
| $\Delta\text{LEC}_{\text{GLU\&GABA}}$ | fam | >0.999 | >0.999 | 0.060 | 0.028 | | |
|  | nov | 0.833 | 0.592 | 0.848 | 0.709 | 0.637 |  |

| dendrites | | control | | $\Delta\text{LEC}_{\text{GLU}}$ | | $\Delta\text{LEC}_{\text{GLU\&GABA}}$ | |
| --- | --- | --- | --- | --- | --- | --- | --- |
|  |  | fam | nov | fam | nov | fam | nov |
| control | fam |  |  |  |  |  |  |
|  | nov | 0.999 |  |  |  |  |  |
| $\Delta\text{LEC}_{\text{GLU}}$ | fam | 0.104 | 0.028 | | | | |
|  | nov | 0.296 | 0.107 | 0.953 |  |  |  |
| $\Delta\text{LEC}_{\text{GLU\&GABA}}$ | fam | 0.988 | >0.999 | <0.001 | 0.005 | | |
|  | nov | 0.985 | 0.879 | 0.080 | 0.350 | 0.557 |  |

**Table S2, related to Figure 7.** GOL non-tuned ROIs event rate p-values (Dunn-Holland-Wolfe *post hoc* test)

| somas | | control | | | $\Delta\text{LEC}_{\text{GLU}}$ | | | $\Delta\text{LEC}_{\text{GLU\&GABA}}$ | | |
| --- | --- | --- | --- | --- | --- | --- | --- | --- | --- | --- |
|  |  | fam | int | nov | fam | int | nov | fam | int | nov |
| control | fam |  |  |  |  |  |  |  |  |  |
|  | int | >0.999 |  |  |  |  |  |  |  |  |
|  | nov | >0.999 | >0.999 |  |  |  |  |  |  |  |
| $\Delta\text{LEC}_{\text{GLU}}$ | fam | 0.025 | 0.038 | 0.020 | | | | | | |
|  | int | 0.191 | 0.246 | 0.172 | >0.999 |  |  |  |  |  |
|  | nov | 0.189 | 0.247 | 0.167 | 0.999 | >0.999 |  |  |  |  |
| $\Delta\text{LEC}_{\text{GLU\&GABA}}$ | fam | 0.994 | 0.989 | 0.994 | 0.002 | 0.028 | 0.024 | | | |
|  | int | >0.999 | >0.999 | 0.999 | 0.168 | 0.566 | 0.591 | 0.888 |  |  |
|  | nov | >0.999 | >0.999 | >0.999 | 0.026 | 0.189 | 0.188 | 0.997 | 0.999 |  |

| dendrites | | control | | | $\Delta\text{LEC}_{\text{GLU}}$ | | | $\Delta\text{LEC}_{\text{GLU}\&\text{GABA}}$ | | |
| --- | --- | --- | --- | --- | --- | --- | --- | --- | --- | --- |
|  |  | fam | int | nov | fam | int | nov | fam | int | nov |
| control | fam |  |  |  |  |  |  |  |  |  |
|  | int | 0.175 |  |  |  |  |  |  |  |  |
|  | nov | 0.626 | 0.997 |  |  |  |  |  |  |  |
| $\Delta\text{LEC}_{\text{GLU}}$ | fam | <0.001 | 0.001 | <0.001 | | | | | | |
|  | int | <0.001 | 0.087 | 0.006 | 0.900 |  |  |  |  |  |
|  | nov | <0.001 | 0.094 | 0.006 | 0.821 | >0.999 |  |  |  |  |
| $\Delta\text{LEC}_{\text{GLU}\&\text{GABA}}$ | fam | >0.999 | 0.159 | 0.600 | <0.001 | <0.001 | <0.001 | | | |
|  | int | 0.999 | 0.604 | 0.965 | <0.001 | <0.001 | <0.001 | 0.998 |  |  |
|  | nov | >0.999 | 0.443 | 0.910 | <0.001 | <0.001 | <0.001 | >0.999 | >0.999 |  |

**Table S3, related to Figure 7.** GOL tuned ROIs event rate p-values (Dunn-Holland-Wolfe *post hoc* test)

| somas | | control | | | $\Delta\text{LEC}_{\text{GLU}}$ | | | $\Delta\text{LEC}_{\text{GLU\&GABA}}$ | | |
| --- | --- | --- | --- | --- | --- | --- | --- | --- | --- | --- |
|  |  | fam | int | nov | fam | int | nov | fam | int | nov |
| control | fam |  |  |  |  |  |  |  |  |  |
|  | int | 0.994 |  |  |  |  |  |  |  |  |
|  | nov | 0.997 | >0.999 |  |  |  |  |  |  |  |
| $\Delta\text{LEC}_{\text{GLU}}$ | fam | 0.999 | >0.999 | >0.999 | | | | | | |
|  | int | >0.999 | 0.993 | 0.997 | 0.998 |  |  |  |  |  |
|  | nov | 0.996 | >0.999 | >0.999 | >0.999 | 0.995 |  |  |  |  |
| $\Delta\text{LEC}_{\text{GLU\&GABA}}$ | fam | 0.974 | 0.557 | 0.768 | 0.560 | 0.941 | 0.524 | | | |
|  | int | >0.999 | 0.949 | 0.979 | 0.973 | >0.999 | 0.952 | 0.995 |  |  |
|  | nov | >0.999 | 0.999 | >0.999 | >0.999 | >0.999 | >0.999 | 0.907 | >0.999 |  |

| dendrites | | control | | | $\Delta\text{LEC}_{\text{GLU}}$ | | | $\Delta\text{LEC}_{\text{GLU}\&\text{GABA}}$ | | |
| --- | --- | --- | --- | --- | --- | --- | --- | --- | --- | --- |
|  |  | fam | int | nov | fam | int | nov | fam | int | nov |
| control | fam |  |  |  |  |  |  |  |  |  |
|  | int | >0.999 |  |  |  |  |  |  |  |  |
|  | nov | 0.827 | 0.927 |  |  |  |  |  |  |  |
| $\Delta\text{LEC}_{\text{GLU}}$ | fam | 0.726 | 0.905 | >0.999 | | | | | | |
|  | int | >0.999 | >0.999 | 0.828 | 0.687 |  |  |  |  |  |
|  | nov | 0.998 | >0.999 | 0.986 | 0.989 | 0.998 |  |  |  |  |
| $\Delta\text{LEC}_{\text{GLU}\&\text{GABA}}$ | fam | 0.996 | 0.976 | 0.386 | 0.129 | 0.984 | 0.766 | | | |
|  | int | >0.999 | 0.997 | 0.536 | 0.257 | 0.999 | 0.912 | >0.999 |  |  |
|  | nov | >0.999 | >0.999 | 0.703 | 0.538 | >0.999 | 0.980 | >0.999 | >0.999 |  |

## Notes

### Competing Interest Statement

The authors have declared no competing interest.

